# Critical role for cold shock protein YB-1 in cytokinesis

**DOI:** 10.1101/2020.03.18.997817

**Authors:** Sunali Mehta, Michael Algie, Tariq Al-Jabri, Cushla McKinney, Srinivasaraghavan Kannan, Chandra S Verma, Weini Ma, Jessie Zhang, Tara K. Bartolec, V. Pragathi Masamsetti, Kim Parker, Luke Henderson, Maree L Gould, Puja Bhatia, Rhodri Harfoot, Megan Chircop, Torsten Kleffmann, Scott B Cohen, Adele G Woolley, Anthony J Cesare, Antony Braithwaite

**Affiliations:** Department of Pathology, University of Otago, Dunedin, New Zealand; Centre for Protein Research, Department of Biochemistry, University of Otago, Dunedin, New Zealand; Maurice Wilkins Centre for Biodiscovery, University of Otago, Dunedin, New Zealand; Children’s Medical Research Institute, University of Sydney, Westmead, New South Wales 2145, Australia; Bioinformatics Institute (A*STAR), 30 Biopolis Street, 07-01 Matrix, Singapore 138671; Malaghan Institute of Medical Research, Wellington, New Zealand

## Abstract

High levels of the cold shock protein Y-box-binding protein-1, YB-1, are tightly correlated with increased cell proliferation and cancer progression. However, the precise mechanism by which YB-1 regulates proliferation is unknown. Here, we found that YB-1 depletion in several cell lines resulted in cytokinesis failure, multinucleation and an increase in G_1_ transit time. Rescue experiments indicated that YB-1 was required for completion of cytokinesis. Using confocal imaging of cells undergoing cytokinesis both *in vitro* and in zebrafish embryos, we found that YB-1 was critical for microtubule organization during cytokinesis. Using mass spectrometry we identified multiple novel phosphorylation sites on YB-1. We show that phosphorylation of YB-1 at multiple serine residues was essential for its function during cytokinesis. Using atomistic modelling we show how multiple phosphorylations alter YB-1 conformation, allowing it to interact with protein partners. Our results establish phosphorylated YB-1 as a critical regulator of cytokinesis, defining for the first time precisely how YB-1 regulates cell division.

**SUMMARY:** Y-box-binding protein-1, YB-1, is essential for cell division, but it is not clear how it functions. Using live imaging and confocal microscopy we show that YB-1 functions only in the last step of division, specifically being required to initiate cytokinesis.

## INTRODUCTION

Y-box binding protein 1 (YB-1) is a multifunctional member of the cold-shock protein superfamily that plays an important role during development and cancer ^1,2^. High levels of *YBX1* mRNA or protein are tightly associated with patient relapse and poor prognosis in multiple cancer types ^3-5^ [reviewed in ^1^]. Analysis of The Cancer Genome Atlas ^6^ further identified that tumors presenting with the highest levels of *YBX1* mRNA and protein were significantly associated with poor patient outcomes (our analysis, Supplementary Fig 1). *In vivo*, YB-1 is required for continued cell proliferation [reviewed in ^1^] and YB-1 overexpression in transgenic mice led to the development of invasive breast cancer in all instances ^7^. In addition, reducing YB-1 levels in tumor xenograft models of breast, brain, lung and pharyngeal cancers inhibited cell proliferation ^4,8,9^. YB-1 has been implicated in regulating many genes involved in proliferation and survival [reviewed in ^1^], including the 70-gene breast cancer “signature” ^10^ and the E2F family gene cluster ^4^. YB-1 has also been shown to disable the p53 pathway to allow continued cellular propagation despite genomic insult ^11^. In addition to these studies, overexpression of YB-1 has been shown to result in cell division errors ^50^. Consistent with damage tolerance, YB-1 also promotes chemotherapy resistance ^12,13^.

In addition to cancer YB-1 is known to be important during development. Specifically, homozygous deletion of YB-1 in mice results in embryonic lethality due to impaired cell proliferation ^14,15^. Knockdown of *ybx1* in zebrafish embryos results in epiboly failure, leading to defects in cell proliferation, pronounced morphological defects and developmental arrest ^16^. Thus, YB-1 is intimately linked with cell proliferation and development.

Several lines of evidence suggest that YB-1 functions in the G_1_ phase of the cell cycle. YB-1 regulates cyclin A and cyclin B1 transcription at the G_1_/S phase border ^17^ and depletion of YB-1 reduced expression of various cyclins and cyclin dependent kinases and increased expression of checkpoint proteins p21 and p16 ^18^. Conversely, ectopic expression of YB-1 increased expression of cyclin D1 and cyclin E ^19^. In contrast, a role for an N-terminal 77 amino acid domain of YB-1 has been implicated in regulating progression of the G_2_/M phases of the cell cycle ^20^ and other studies have demonstrated that inhibiting YB-1 caused an arrest at G_2_/M ^21,22^. Furthermore, prolonged exposure of breast cancer cells to YB-1 led to cytokinesis failure and slippage through the G_1_/S border ^23^. Thus, it is unclear from these reports whether YB-1 functions in one or more cell cycle phases.

To address this issue, we used live single cell imaging combined with high end confocal microscopy to define precisely how YB-1 regulates the cell cycle. Our results show that YB-1 facilitates the assembly of microtubules and specification of the cleavage plane in the equatorial region during cytokinesis, the last step in cell division, without affecting other cell cycle phases. This establishes YB-1 as a critical regulator of cytokinesis.

## RESULTS

### YB-1 depletion causes cytokinesis failure, multinucleation and accumulation in G_1_

To investigate YB-1 function in the cell cycle, live cells were tracked using time lapse imaging after YB-1 depletion. Five cell lines were used, including 4 transduced with the FUCCI (F) cell cycle reporter constructs ^24^: A549/F lung cancer cells with wild type (wt)*TP53* ^25^; HT1080/F fibrosarcoma cells with wt*TP53* ^26^ and the variant HT1080-6TG (6TG/F) with mutant *TP53* ^27^; the spontaneously immortalized IIICF/c/F skin fibroblast line from a Li Fraumeni patient ^28^; and MDA-MB-231 breast cancer cells with mutant *TP53*. Cells were transfected with control (si-Ctrl) or YB-1 siRNA (si-YB-1) and western blotted to confirm YB-1 depletion (Supplementary Fig 2a). Live-cell imaging was started 24 hours post siRNA transfection with images collected using differential interference contrast (DIC) bright field and fluorescent microscopy every 6 minutes for 60 hours. In all cell lines tested, YB-1 depletion induced a striking phenotype of disrupted cytokinesis (Fig 1 a-c). Live cell imaging showed that 50-80% of mitoses in YB-1 siRNA treated cultures failed to complete cytokinesis, compared to approximately 10% in the controls (Fig 1b, c). Transfection of an shRNA targeting endogenous YB-1 3’-UTR (sh-YB-1) also depleted YB-1 and induced cytokinesis failure (Fig 1b). This phenotype was rescued by co-transfecting a YB-1^EBFP2^ expressing construct (Fig 1b), confirming that cytokinesis failure is due specifically to loss of YB-1. As expected, cytokinesis failure resulted in significant accumulation of multinucleated cells over time (Fig 1d, Supplementary Videos 1-2) and overexpression of YB-1^EBFP2^ along with sh-YB-1 significantly decreased the number of multinucleated cells (Fig 1d). Finally, depletion of YB-1 resulted in reduced cell proliferation (Supplementary Fig S3a). Of interest, YB-1 depletion also increased the G_1_ duration of the nuclei in (multinucleated) A549/F and HT1080/F cells, but not in the nuceli of 6TG/F or IIICF/c/F cells (Fig 1e-f) and co-expression of YB-1^EBFP2^ along with YB1 sh-RNA rescued the increased G_1_ transit timeduration (Fig 1e). To test if this was p53 dependent, A549/F cells were co-transfected with siRNAs against p53 (si-p53) and YB-1. Co-depletion of YB-1 and p53 (Supplementary Fig 2a and 2c) significantly decreased G_1_ duration (Fig 1e). Collectively, the above results show that YB-1 regulates cytokinesis and in turn cell division independently of p53, but causes an increase in G_1_ duration of the daughter nuclei, that is p53 dependent.

**Figure 1:**
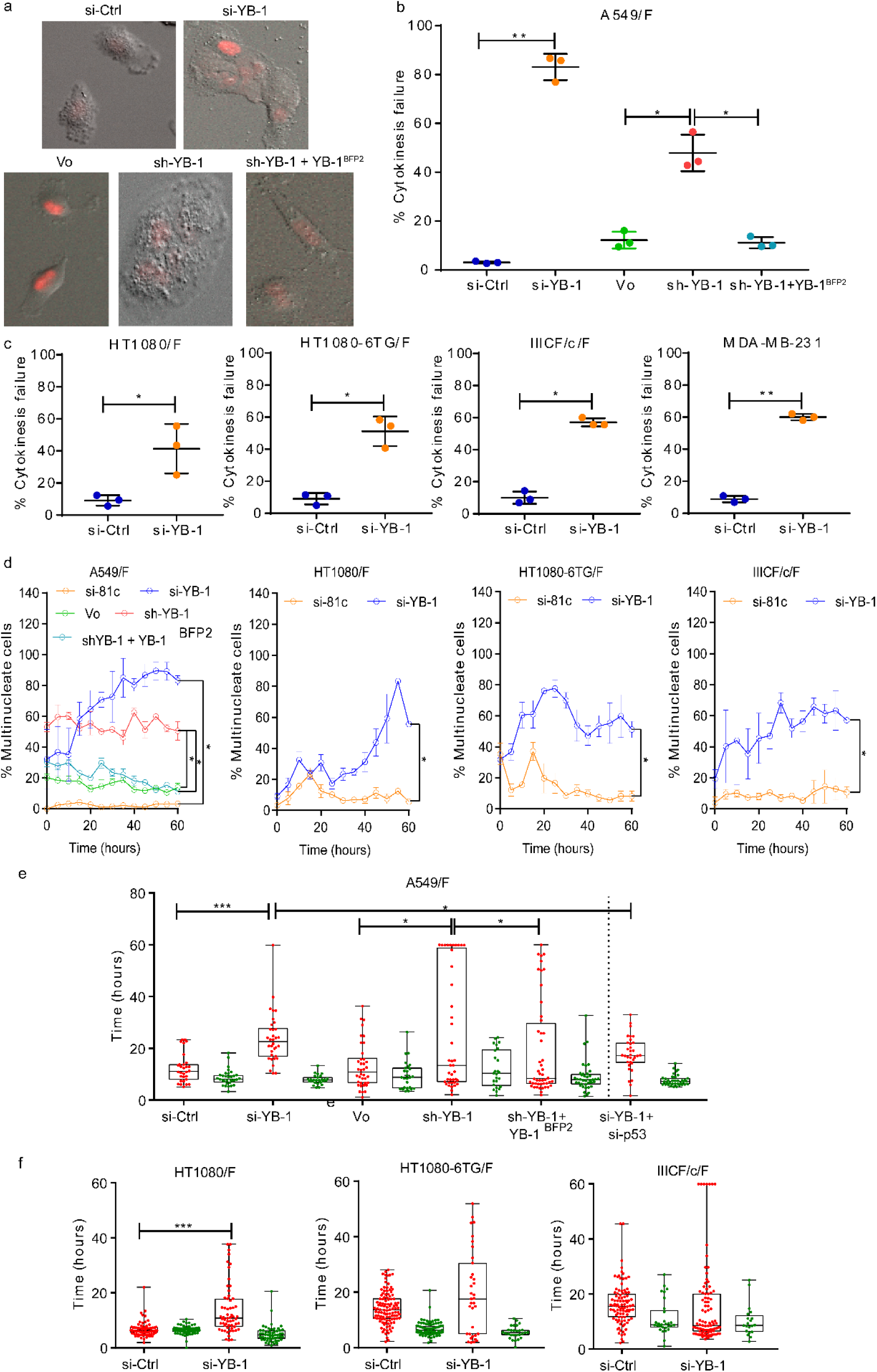
YB-1 depletion inhibits cytokinesis resulting in multinucleation and an increase in G_1_ transit time. **a.** Examples of A549/F cells treated with si-Ctrl, si-YB-1, Vo, sh-YB-1 or sh-YB-1+YB-1^EBFP2^. **b.** Percentage of A549/F cells that fail cytokinesis treated with si-Ctrl, si-YB-1, sh-YB-1, or sh-YB-1 + shRNA-resistant YB-1^EBFP2^. **c.** Percentage of cells that fail cytokinesis in HT1080/F, HT1080-6TG/F, IIICF/c/F and MDA-MB-231 cells treated with si-Ctrl and si-YB-1. **b-c.** n ≥ 100 cells from three independent image fields from one experiment. Dot plots with lines shown mean ± s.d. and one tailed student’s t test, * p<0.05, ** p<0.01, ** p<0.001. **d.** Percentage of multinucleated A549/F cells in cultures treated with si-Ctrl, si-YB-1, Vo, sh-YB-1 and sh-YB-1+YB-1^EBFP2^, 60 hours post 24h transfection and percentage of multinucleated HT1080/F, HT1080-6TG/F and IIICF/c/F cells treated with either si-Ctrl or si-YB-1 80 hours post transfection. Each point represents the mean ± s.e.m for 3 independent imaging fields from one experiment. Significance was determined using multiple t-tests with FDR correction, * p < 0.05. **e.** Distribution of the G_1_ and S-G_2_-M phases of A549/F cells treated with either si-Ctrl, si-YB-1, si-YB-1+si-p53, Vo, sh-YB-1 and sh-YB-1+ YB-1^EBFP2^. **f.** Distribution of G_1_ and S-G_2_-M phases of HT1080/F, HT1080-6TG/F and IIICF/c/F cells treated with either si-Ctrl or si-YB-1 **e-f.** The line in the middle of each box represents the median, the top and bottom outlines of the box represent the first and third quartiles. Significance was determined using Mann-Whitney U test, * p < 0.05, ** p < 0.01, *** p < 0.001 and **** p < 0.0001.

### YB-1 is required for initiation of cleavage furrow formation

Cytokinesis is the final step during cell division required for formation of two daughter cells ^29^. Cytokinesis begins after chromosome segregation, followed by defining the cleavage plane resulting in formation of a cleavage furrow, ingression and abscission, which ends with the physical separation of the daughter cells ^29^. Cleavage furrow ingression is controlled by proteins involved in regulating spindle assembly dynamics, which include microtubule associated proteins (α-tubulin), the centraspindlin complex, and the chromosome passenger complex (CPC), comprised of Survivin, Borealin, Inner Centromere Protein (INCENP), and Aurora B kinase (AURKB) ^30^. CPC localization at the equatorial cortex is required for sustained cleavage furrow ingression ^30,31^. One of the key requirements for completion of cytokinesis is the assembly of the microtubules, specification of the cleavage plane and timely localization of the chromosome passenger complex (CPC) to the cleavage furrow ^29,30^. Defects in any of these processes will result in cytokinesis failure.

To define which cytokinesis stage is affected by YB-1 depletion we analyzed our live cell imaging data, focusing on phenotypes during late mitosis and cytokinesis (Supplementary videos 1-2). DIC live cell imaging revealed that cells with reduced YB-1 form a ‘twisted’ cleavage furrow (Fig 2a, see arrows) at a significantly higher frequency than those treated with the si-Ctrl (Fig 2b), thus preventing ingression and consequently cytokinesis. This suggests that YB-1 is required for formation of the cleavage furrow. To test this, we first used confocal microscopy to determine whether YB-1 is localised at or near the cleavage furrow. Results (Fig 2c) show that YB-1 is enriched at the mid-zone and appears to intercalate within AURKB. To understand better how YB-1 functions in cytokinesis, additional confocal microscopy was carried out to include proteins known to localise at the cleavage furrow; α-tubulin, AURKB, CAPZA1 ^32^ and MSN ^33,34^. In parallel, we did a similar experiment in YB-1 depleted A549 cells. Samples were collected at 48 hours after treating cells with either si-Ctrl or si-YB-1. In control cells, (Fig 2d-e) YB-1 was again enriched at the mid-zone localising with α-tubulin and with AURKB, CAPZA1 and MSN. In YB-1 depleted cells all these proteins failed to localise to the midzone, remaining dispersed throughout the daughter nuclei (Fig 2d-e). This was not due to reduced protein levels (Supplementary Fig 2d). Thus, it appears that YB-1 is required for cleavage furrow formation.

**Figure 2:**
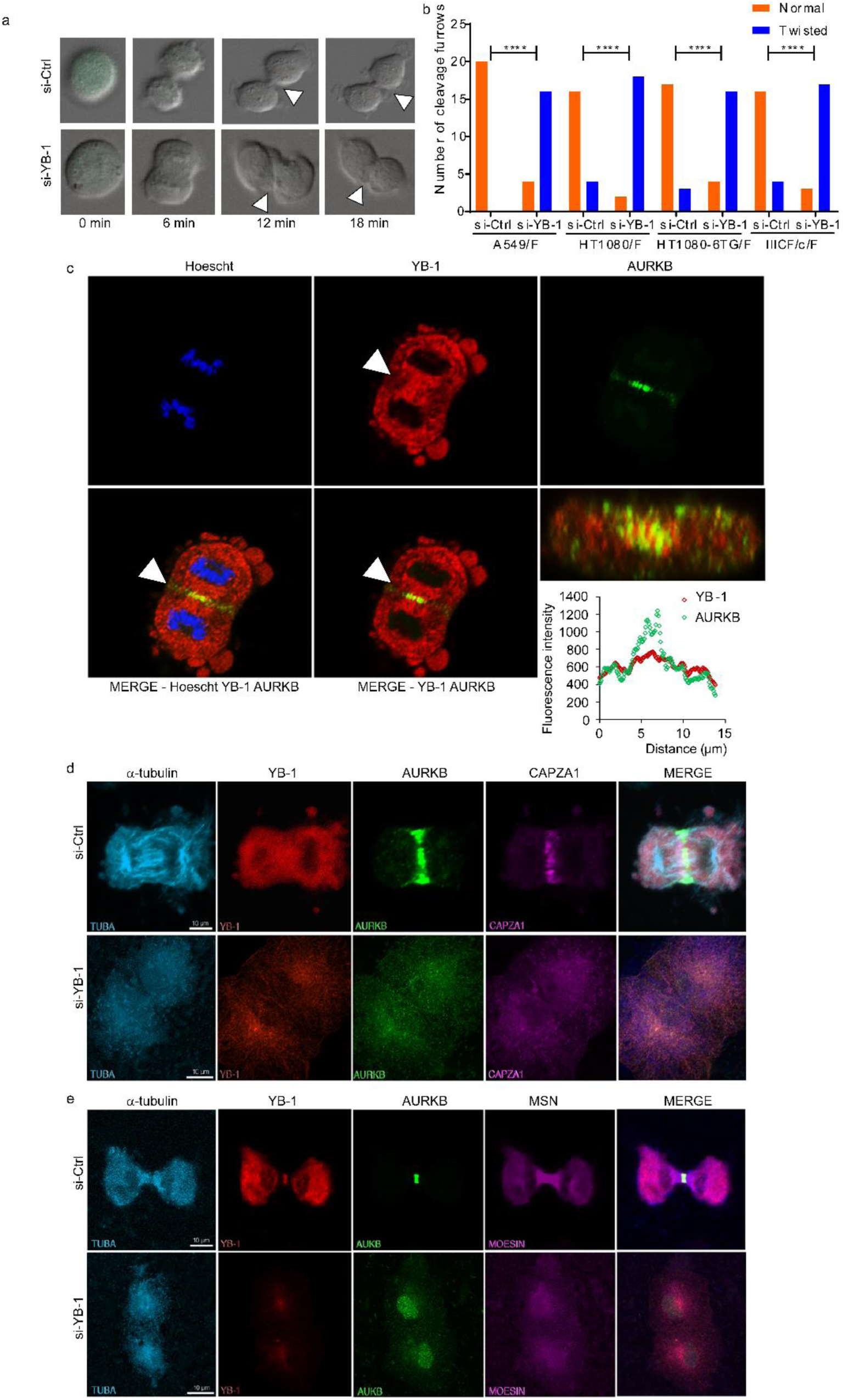
YB-1 depletion results in the failure of Chromosome Passenger Complex (CPC) recruitment to the cleavage furrow. **a.** Examples from live cell imaging of an A549/F cell treated with si-Ctrl and forming a stable cleavage furrow (left panel) or si-YB-1 and forming a twisted cleavage furrow (right panel). **b.** Quantitation of normal versus twisted cleavage furrows in A549/F, HT1080/F, 6TG/F and IIICF/c/F cells treated with either control siRNA (si-Ctrl) or targeting YB-1 (si-YB-1). Twenty cells were counted for each condition. Significance was determined using Chi-Square and Fishers Exact test. p < 0.05 was considered to be significant. **** p < 0.0001. **c.** Localisation of YB-1 to the cleavage furrow in A549 cells. **d - e.** A549 cells were treated with either an si-Ctrl (top row) or an si-YB-1 (bottom row) and subsequently immunostained with antibodies against (from left to right) α-tubulin (cyan), YB-1 (red), AURKB (green), CAPZA1 or MSN (magenta) together with a merged image of all four. **d.** Accumulation of CAPZA1 to the cleavage furrow and **e.** accumulation of MSN to the cleavage furrow.

Formation of the cleavage furrow requires establishment of the cleavage plane during late metaphase, together with timely localisation of the CPC and organisation of the microtubule fibers at the midzone ^30,31^. However, the mechanism that is responsible for establishment of the cleavage plane is unknown ^29^. To test if YB-1 is required for establishment of the cleavage plane, we synchronized A549 cells at G_1_ using a double thymidine block, released the block and once the cells had reached mitosis (∼7 hours later) fixed them sequentially at 5 minutes intervals for up to 90 minutes to capture those progressing towards cytokinesis. Cells were then stained for YB-1, α-tubulin, and AURKB and imaged using confocal microscopy. Results show that during prophase YB-1 appears as a diffuse ring around AURKB and localizes with α-tubulin at the midzone (Figure 3 – top row). During prometaphase, YB-1 is enriched in the region of the microtubules and again surrounds AURKB (Figure 3 – row 2). During metaphase AURKB is tightly localized to the midzone, whilst YB-1 is both disperse and localized with α-tubulin at the spindle poles, flanking the cleavage plane (Figure 3 – row 3). Finally, during anaphase, AURKB defines the contractile ring and YB-1 continues to associate with α-tubulin (Figure 3 – bottom row). Taken together, these data suggest that YB-1 is required for the organization of microtubules needed for CPC attachment in the equatorial zone to define the cleavage plane.

**Figure 3:**
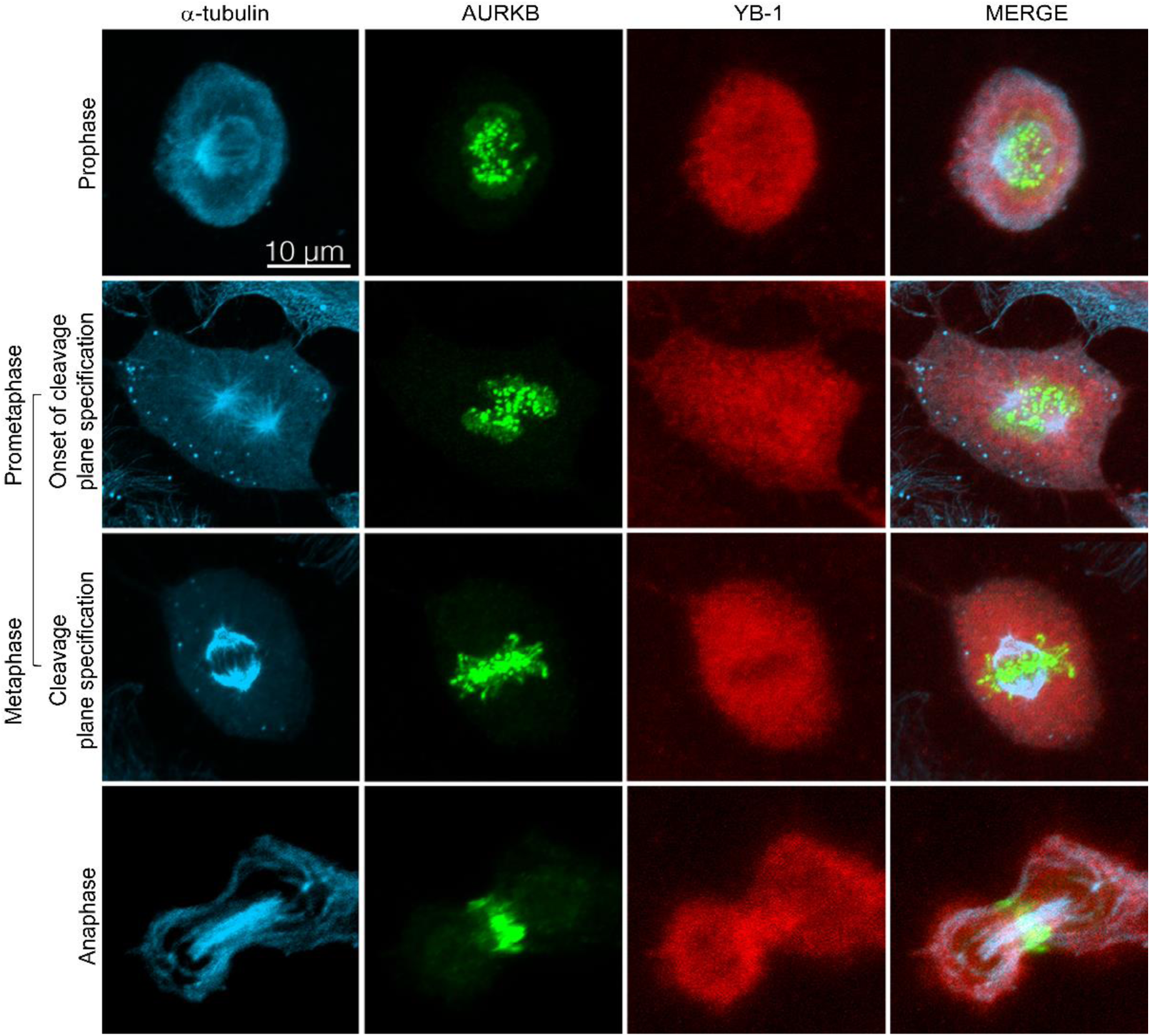
YB-1 initiates formation of the cleavage furrow by defining the plane of division. Examples of synchronized A549 cells enriched for cytokinesis, immunostained with antibodies against (from left to right) α-tubulin (cyan), AURKB (green), YB-1 (red), followed by a merged image of all three. The top row shows the presence of YB-1 at the mid-zone, assembly of the microtubule structure (α-tubulin) and a diffuse band of AURKB around the mid-zone. The second row shows cells in prometaphase, with YB-1 at the mid-zone localizing with α-tubulin and the localization of AURKB to the mid-zone. The third row shows cells in metaphase, where YB-1 is both disperse and localized with α-tubulin fibres but excluded from the mid-zone, and AURKB is localized in the mid-zone. The fourth row shows cells in anaphase, in which YB-1 is localized with α-tubulin now incorporated into the contractile ring and AURKB is still localized in the mid-zone.

### YB-1 depletion causes cytokinesis failure in zebrafish embryos resulting in developmental defects

Zebrafish ybx1 protein has been shown to be essential for development and either mutations or depletion of zebrafish y*bx1* resulted in failure to initiate epiboly, reduced cell proliferation and eventually developmental arrest ^16,35-37^. To determine if the cause of the developmental defects could be due to cytokinesis failure, we injected morpholino-oligonucleotides (MO) targeted to *ybx1* or a non-targeting control, at the one or two cell stage and characterized the effects using DIC and confocal microscopy. DIC images of embryos treated with MO_*ybx1* showed failure to initiate epiboly due to impaired cell proliferation compared to the control (Fig 4a) as had been previously reported by ^16^. Moreover, MO_ybx1 treated embryos resulted in developmentally defective fish 48h post fertilization (Fig 4a, Supplementary Fig 2e). Next, to establish if failure of epiboly initiation in *ybx1*-depleted embryos is due to a disruption in the organization of microtubules, we performed whole-mount immunofluorescence for ybx1 and α-tubulin and images were acquired by confocal microscopy. Results show that in MO_*ybx1* treated embryos there was a failure in microtubule organization and assembly compared to the controls. This leads to a failure in cleavage plane specification (Fig 4b). These results suggest that YB-1 is essential for regulating the process of cytokinesis during development.

**Figure 4:**
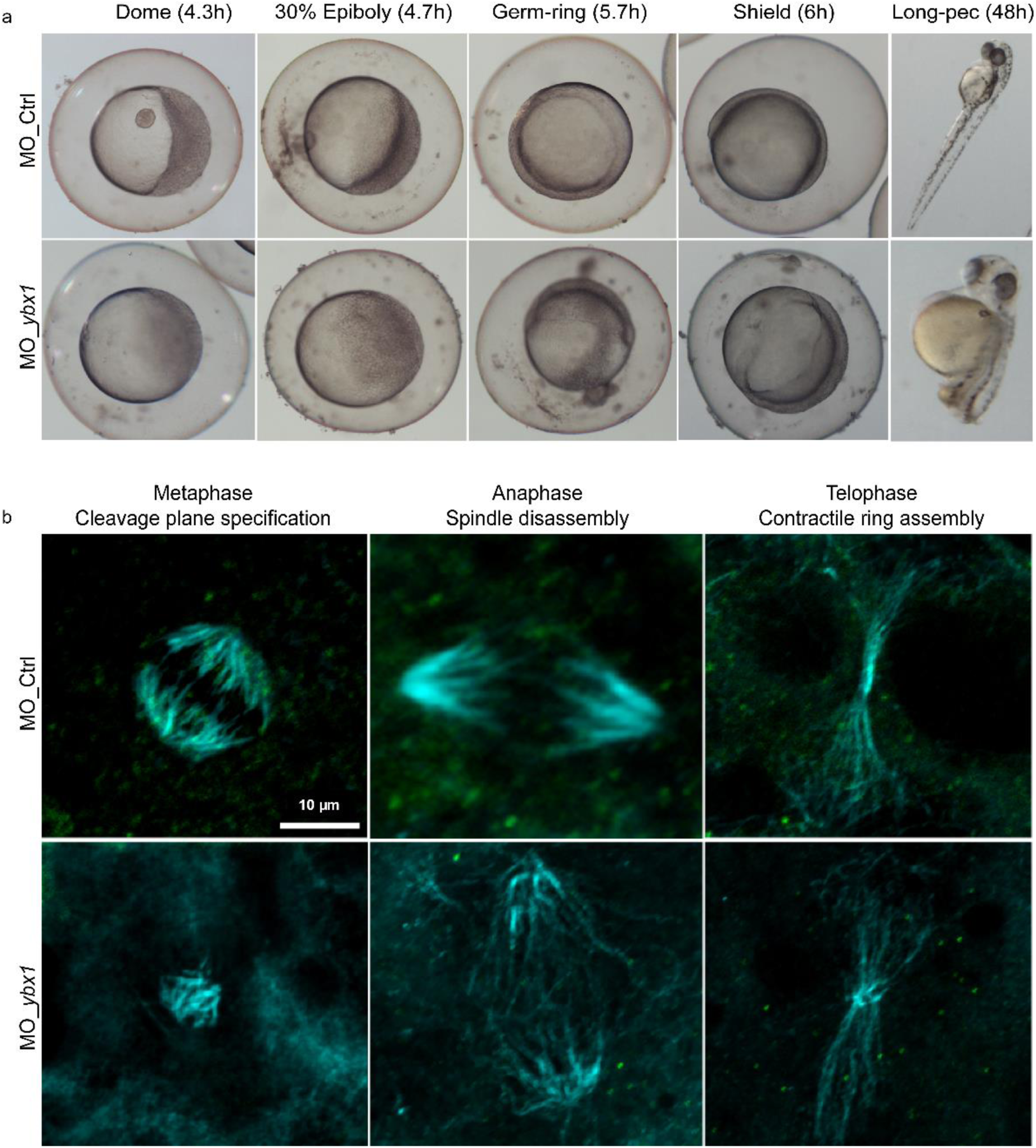
YB-1 depletion results in developmental defects due to cytokinesis failure in zebrafish embryos. **a.** Examples of zebrafish embryos at 4.3, 4.7, 5.7, 6 and 48 hours post fertilization treated 4ng of either control morpholino (MO_Ctrl, top panel) or a morpholino targeting *ybx1* (MO_*ybx1*, bottom panel). **b.** Examples of zebrafish embryonic cells that have been treated with 4ng of MO_Ctrl (top panel) or MO_*ybx1* (bottom panel) 5.7 – 6h post fertilization undergoing cytokinesis, immunostained with antibodies to YB-1 (green) and α-tubulin (cyan). The top-left panel shows the presence of the YB-1 at the mid-zone localized with organized α-tubulin; the top-middle panel shows the localization of YB-1 with α-tubulin flanking the mid-zone; and the top-right panel shows that YB-1 continues to localize with α-tubulin providing a scaffold for assembly of the contractile ring during telophase. The bottom panel (left to right) shows depletion of *ybx1*, lack of correct α-tubulin organization, and absence of the cleavage plane.

### p90 Ribosomal-S6 Kinase (RSK) phosphorylates YB-1 and is required for cytokinesis

YB-1 function is thought to be regulated through phosphorylation ^38^. Inhibiting YB-1 phosphorylation on serine 102 with a mutation to alanine (S102A) prevented cell proliferation ^39^ and transactivation of various genes including *EGFR* ^40^ and *PIK3CA* ^41^. Protein Kinase B (AKT) ^39^, p90 S6 Ribosomal Kinase (RSK) and Extracellular Signal Regulated Kinase (ERK) have all been implicated in YB-1 S102 phosphorylation, and inhibiting RSK suppressed the ability of YB-1 to promote proliferation ^42^. To identify putative kinases capable of phosphorylating YB-1 at either S102 or at other sites on the protein, we used in silico Group-based Prediction System 2.1 ^43^. This identified four candidate kinases: AKT ^39^, RSK ^42^, Casein kinase 2 (CK2) and AURKB.

To determine whether inhibiting putative YB-1 regulators affected cytokinesis, A549/F cells were treated over several days with solvent (DMSO), or different concentrations of Barasertib (AURKB inhibitor), BI-D1870 (RSK inhibitor), MK-2206HCl (AKT inhibitor) or Silmitasertib (CK2 inhibitor). Cultures were then visualized using an IncuCyte live imaging system and cell proliferation analyzed to determine a non-cytotoxic dose for each compound (Supplementary Fig S3b). A549/F cells were then treated with the various inhibitors at the determined concentrations and live cell imaging was performed as described above, recording images every 6 minutes for 46 hours. Consistent with their known function in promoting cytokinesis, inhibiting AURKB ^44^ or RSK ^45^ resulted in cytokinesis failure and multi-nucleated cells (Fig 5a-b) as was observed with YB-1 depletion. RSK inhibition with BI-D1870 also conferred a delay in cleavage furrow formation (Fig 5a). All four kinase inhibitors resulted in increased G_1_ duration (Fig 5c, left panel), while RSK inhibition also resulted in an approximate three-fold increase in S/G_2_/M duration (Fig 5c, right panel). A small but significant S/G_2_/M duration increase was also observed with the CK2 inhibitor.

**Figure 5:**
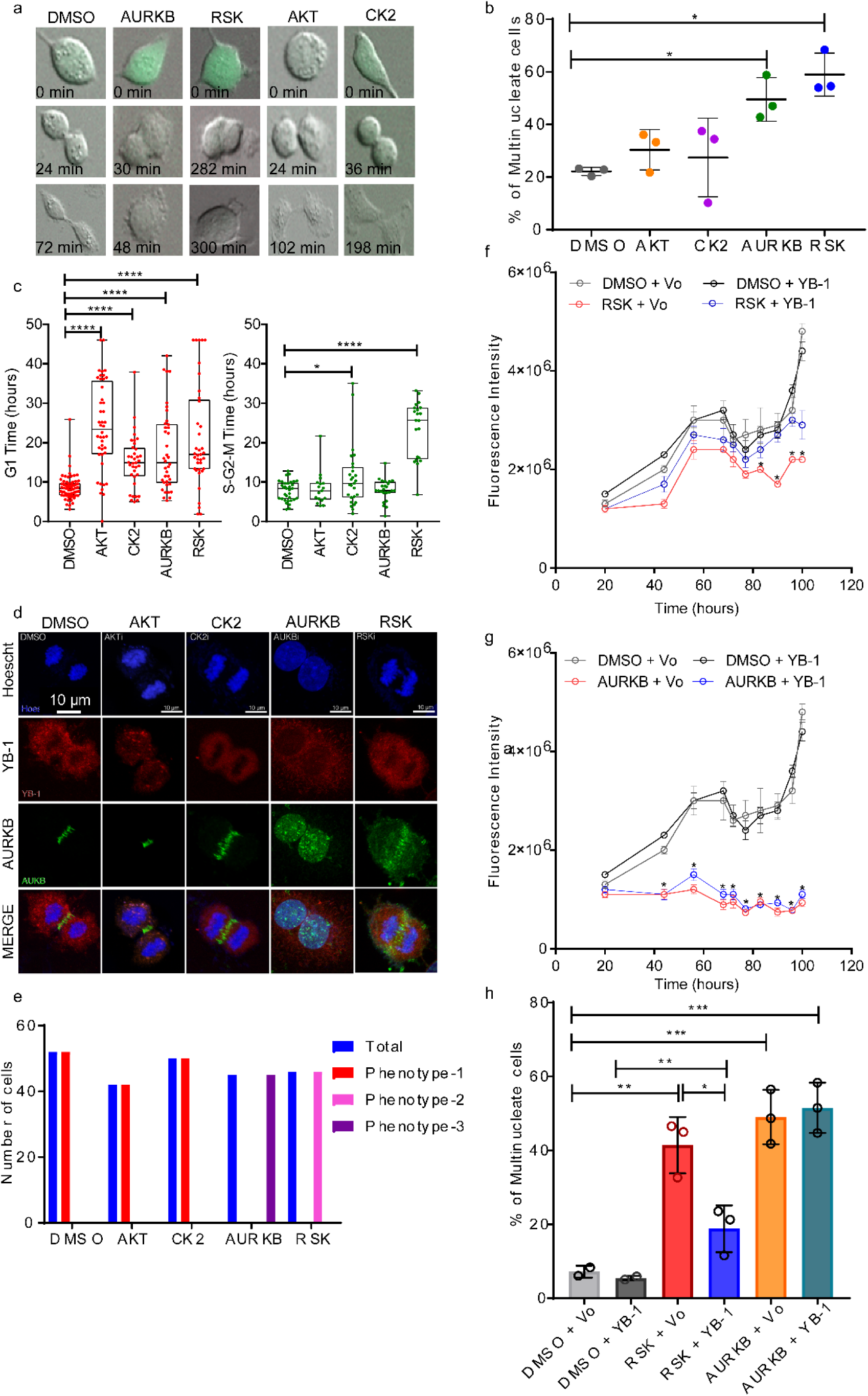
RSK inhibition phenocopies YB-1 depletion. Results from cells treated with solvent control (DMSO), Barasertib-1µM (AURKB), BI-D1870-5µM (RSK), MK-2206HCl-10µM (AKT) or Silmitasertib-10µM (CK2). **a.** Examples of failed cytokinesis after treating A549/F cells. **b.** Percentage of multinucleate A549/F cells 40 hours post treatment. n ≥ 100 cells, 3 independent image fields, one experiment. Dot plots with lines shown mean ± s.d., one tailed student t test, * p<0.05. **c.** Distribution of time in G_1_ (left) and in S-G_2_-M phases (right) of A549/F cells post treatment, Mann-Whitney U test, * p < 0.05 and **** p < 0.0001. **d.** Example of A549 cells post treatment undergoing cytokinesis stained with Hoechst (blue), anti-YB-1 (red) or anti-AURKB (green). **e.** Quantitation of AURKB staining at the cleavage furrow in A549 cells; Phenotype1 – tightly localized, Phenotype 2 – diffuse, Phenotype 3 - diffuse nuclear staining. **f-g**. Shows the amount of SYBR Green I – fluorescence over time in cells transfected with Vo or YB-1^EBFP2^ and treated with DMSO or BI-D1870 (**f)** or Barasertib (**g**). Each point represents the mean ± s.e.m in ≥ 3 replicates, student’s t-tests * p < 0.05. **h.** Percentage of multinucleate cells in A549/F cells treated with 200ng of Vo or YB-1 and either DMSO, BI-D1870 or Barasertib for 40 hours, one-way ANOVA with Tukey’s multiple comparison test. * p < 0.05, ** p < 0.01, *** p < 0.001.

We then assessed how kinase inhibition affected cytokinesis and CPC localization using AURKB as a marker of CPC at the midzone. Control, CK2 and AKT inhibition resulted in normal CPC enrichment at the equatorial cortex (phenotype-1, Fig 5d-e). However, with RSK inhibition, AURKB only partially enriched at the equatorial cortex and was also located at the polar cortex (Phenotype-2, Fig 5d-e). As expected, no cleavage furrow formed in cells treated with Barasertib, where AURKB remained in the nucleus (Phenotype-3, Fig 5d-e). Consistent with RSK-dependent regulation of S102, we observed YB-1 phosphorylation at S102 was reduced in cells when RSK was pharmacologically inhibited with BI-D1870 but not with the AURKB inhibitor Barasertib (Supplementary Fig 2f). Cumulatively, of the kinases tested, only inhibiting RSK reduced YB-1 phosphorylation and phenocopied cytokinesis failure observed with YB-1 knockdown. This suggests RSK regulates YB-1 function during cytokinesis. In agreement, we found that over-expressing YB1^EBFP2^ partially rescued the proliferation and multinucleation phenotypes induced with RSK, but not AURKB inhibition (Fig 5f-h). Also, the phosphorylation defective S102A mutant failed to rescue from the multinucleation phenotype following knockdown of YB-1 (Fig 6b-d).

**Figure 6:**
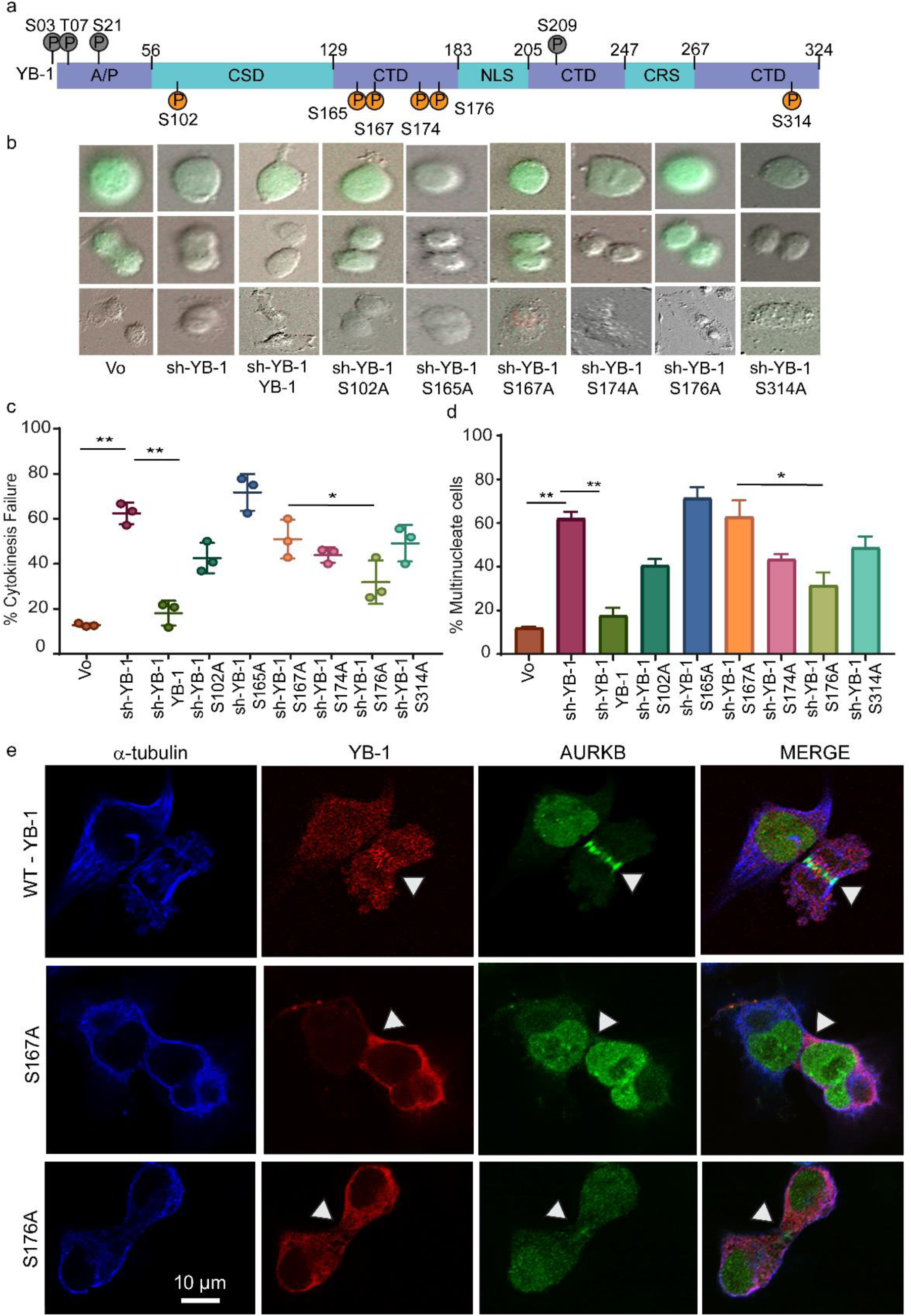
Phosphorylation defective YB-1 mutants cannot rescue cytokinesis failure following YB-1 depletion. **a.** Schematic showing the YB-1 protein and the location of the phosphorylation sites detected in MDA-MB-231, A549 and T47D cells (Common – orange circles; summarized in Supplementary Table 1, and rare – gray circles; summarized in Supplementary Table 2). A/P – Alanine/Proline rich domain; CSD – Cold Shock Domain, CTD – C–terminal Domain, NLS – nuclear localization signal and CRS – cytoplasmic retention signal. **b-e.** Data collected from A549/F cells treated with either Vo or sh-YB-1 in combination with Vo or YB-1^EBFP2^ or the six phosphorylation defective mutants (S102A, S165A, S167A, S174A, S176A and S314A). **b.** Examples of A549/F cells undergoing cytokinesis defects. **c.** Quantitation of the experiment shown in (b). Each point represents the mean ± s.e.m. for ≥ 3 independent image fields from one experiment. Significance was determined using one-way ANOVA followed by Tukeys multiple comparison test. * p < 0.05 **d.** Shows percentage of multinucleated cells 64h post treatment with Vo or sh-YB-1 in combination with Vo or YB-1 or the six phosphorylation defective mutants (S102A, S165A, S167A, S174A, S176A and S314A). Each point represents the mean ± s.e.m for 8 imaging fields from one experiment. Significance was determined using multiple t-tests with FDR correction, * q < 0.05. **e.** Examples of CPC staining in A549 cells after treatment with sh-YB1 in combination with either YB-1, S167A or S176A mutants. Cells are stained (left to right) with antibodies to α-tubulin (blue), YB-1 (red) or AURKB (green) followed by merged images.

### Phosphorylation defective YB-1 mutants fail to complete cytokinesis

In addition to phosphorylation at S102, phosphorylated serines at residue 165 (pS165) and 176 (pS176) were recently identified by mass spectrometry ^46,47^. Mutating these residues impaired the ability of YB-1 to transactivate the NFκB promoter, reduced cell proliferation, and impacted tumor progression ^46,47^. To determine which phosphorylation sites on YB-1 are important for cytokinesis, we immunopurified YB-1 from 3 cell lines (A549 and 2 breast cancer lines; MDA-MB-231 and T47D). YB-1 was then electrophoretically separated by SDS-PAGE, excised, digested with trypsin, and phosphorylated peptides enriched with TiO_2_ were then analyzed by LCMS/MS. We identified 10 phosphorylated residues (Supplementary Fig. 4a and Table 1), 5 of which were identified in all 3 cell lines in multiple repeat experiments, including pS165 and pS176 and novel phosphorylated residues pS167, pS174 and pS314 (Table 1, Fig 6a). Of note, the most frequently studied (pS102) residue appeared to be difficult to detect in LC-MS/MS as it was observed in only 3/8 runs and in only one cell line (Table 1, Fig 6a). Of interest, all the common phosphorylated residues and pS102 are conserved in higher mammalian species and none of these 6 sites is mutated in 7799 tumors across 32 different tumor types analyzed using The Cancer Genome Altas ^6^ data from cBioPortal ^48^. Thus phosphorylation of these residues maybe critical for YB-1 function. All of the above sites except for S176 are predicted to be phosphorylated by RSK using the Group-based Prediction System 2.1. Using targeted mass spectrometry we show that inhibition of RSK with BI-D1870 reduced phosphorylation of YB-1 at S165, S167, S174 and S176, whereas AURKB inhibition with Barasertib only reduced phosphorylation at S165 and S167 (Supplementary Fig 4b).

**Table 1:**
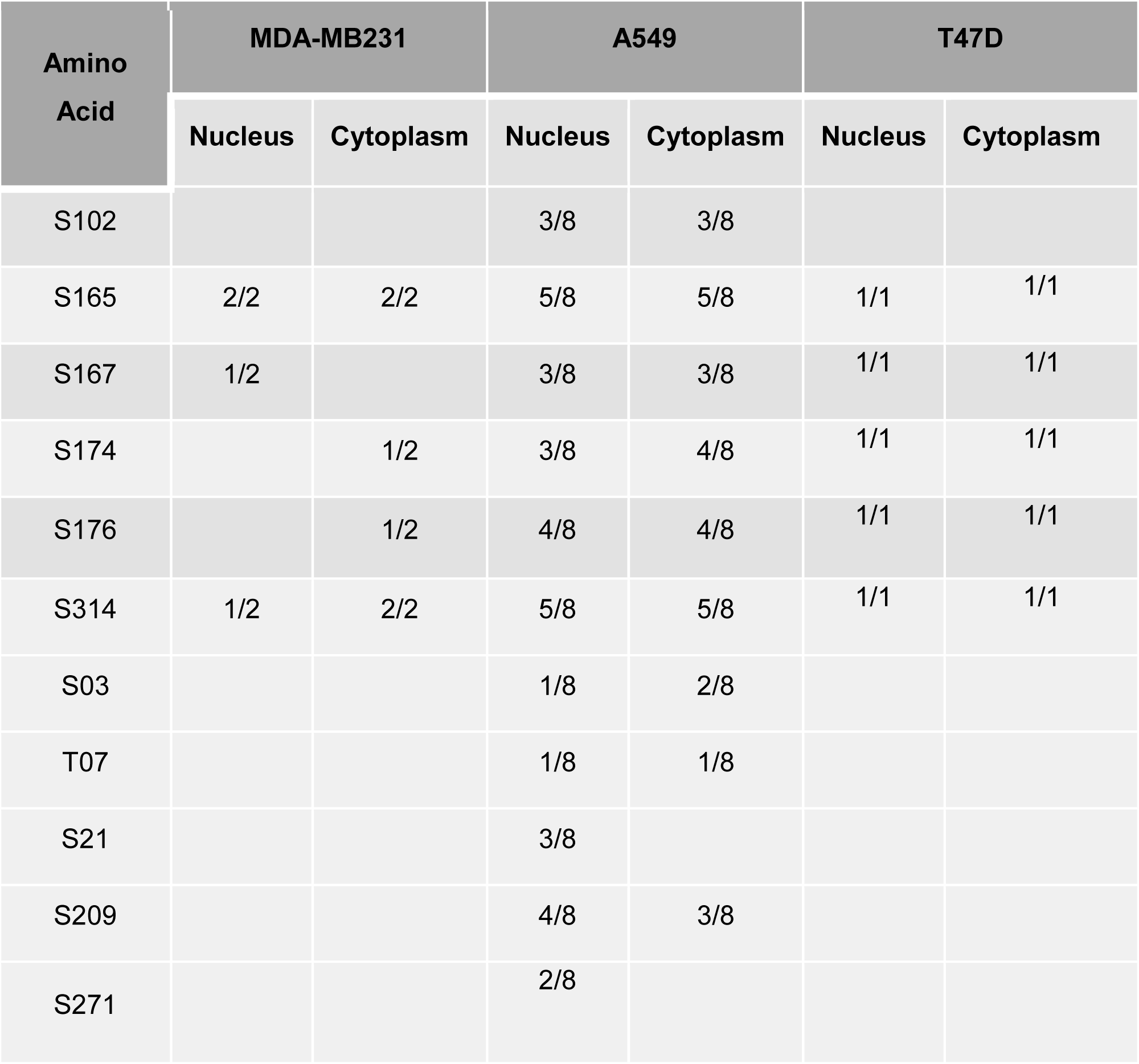
Summary of phosphorylation sites identified by LC-MS/MS in A549, MDA-MB-231 and T47D cells. Numerator – number of times phosphorylation site was detected, Denominator – total number of LC-MS/MS runs.

To determine whether phosphorylation of these conserved residues contributes to YB-1 function in cytokinesis, we created YB-1^EBFP2^ expression constructs with an alanine substitution at each phosphorylated serine. A549/F cells were transfected with sh-YB-1 with or without co-expression of YB-1^EBFP2^ or phospho-defective mutants S102A, S165A, S167A, S174A, S176A and S314A. Cultures were imaged and analyzed as described above. The mean blue fluorescence intensity was used as a marker for expression of the plasmids which showed that all exogenous mutant proteins were expressed at similar levels (Supplementary Fig 4c). Interestingly, all 6 phosphorylation defective mutants failed to rescue the cytokinesis defects induced by depleting endogenous YB-1, although S176A was attenuated (Fig 6b-d). To determine how YB-1 phosphorylation affected CPC recruitment during cytokinesis, we co-expressed sh-YB-1 with S167A or S176A and performed immunofluorescence. Notably cells co-expressing the sh-YB-1 and S167A failed to form the cleavage furrow and AURKB remained primarily dispersed in the nucleus (Fig 6e), while S176A enabled partial cleavage furrow formation and AURKB localization (Fig 6e).

To determine the impact of phosphorylated YB-1 on other cell cycle phases, we quantified G_1_ and S/G_2_/M duration in A549/F cells treated with sh-YB-1 along with YB-1 or phosphorylation defective YB-1 mutants. Cells treated with sh-YB-1 mostly stalled in the G_1_ phase of the cell cycle which was overcome by YB-1^EBFP2^ co-expression, as indicated by alternating red and green (cycling) cells (Figs. 7a and 7c). With the exception of S176A, all mutants were stalled in G_1_ (S Figs. 7b-c). By contrast, ∼30% of cells expressing S176A continued to cycle, consistent with this mutant being attenuated. Finally, with the exception of S165A and S314A, none of the mutants increased the time spent in S-G_2_-M phase of the cell cycle (Figs. 7b and 7d). Collectively these data suggest that phosphorylation on all six YB-1 sites is required to complete cytokinesis and progress through the cell cycle.

**Figure 7:**
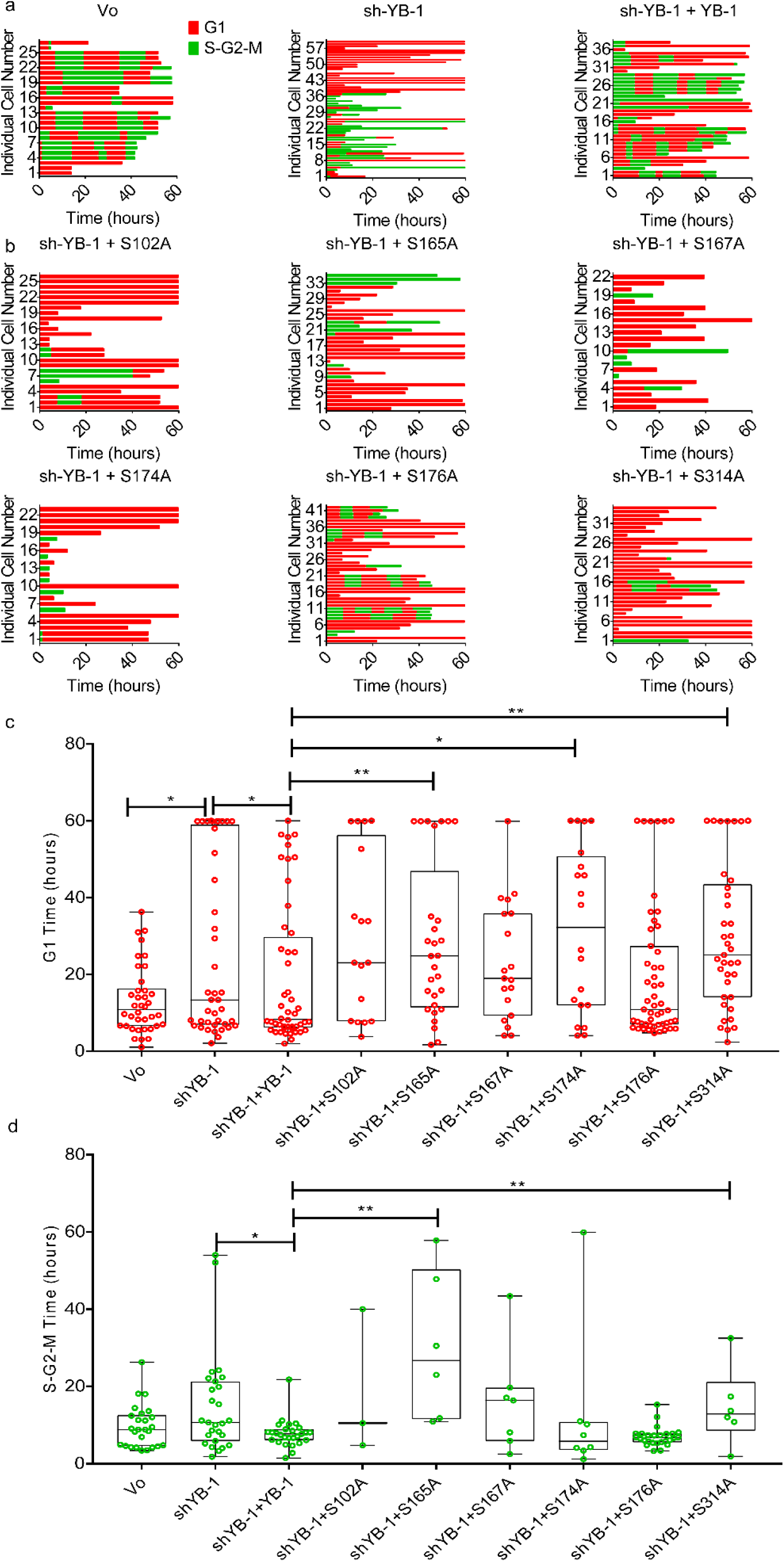
Phosphorylation defective YB-1 mutants do not rescue from cytokinesis failure. **a – b.** Shows the starting phase of the cell cycle for each tracked A549/F cell using time lapse imaging and quantitates the amount of time spent in each phase of the cell cycle and the number of divisions that each cell undergoes. **c.** Quantitation of G1 phase transit time. **d.** Quantitation of S-G2-M phase transit time. **c-d**. Significance was determined using Mann Whitney U test, * p < 0.05, ** p < 0.01, *** p < 0.001 and **** p < 0.0001.

### Cooperative YB-1 phosphorylation on multiple residues alters protein folding

Phosphorylation on multiple residues commonly confers structural rearrangements, regulating protein-protein interactions. To determine how YB-1 phosphorylation affects protein folding, we generated atomic models of YB-1 using the I-TASSER pipeline ^49^. YB-1 protein was modelled using the known structure of the cold shock domain (CSD, residues 50-124) ^50^ with and without phosphorylated residues. The selected template depicts YB-1 to be a disordered protein as has been reported ^51^. Based on the model, there are positive charges including Arginine and Lysine (thick grey lines) both within the CSD and at the N and C termini of YB-1 (Fig 8a). Simulations of molecular dynamics were carried out to determine structural changes resulting from phosphorylation. Results suggest that phosphorylation causes YB-1 to fold such that a negatively charged interior is created, which exposes positively charged residues at the protein surface. This is anticipated to create a positively charged binding patch capable of interacting with negatively charged protein domains or nucleic acids (Fig 8a-c).

**Figure 8:**
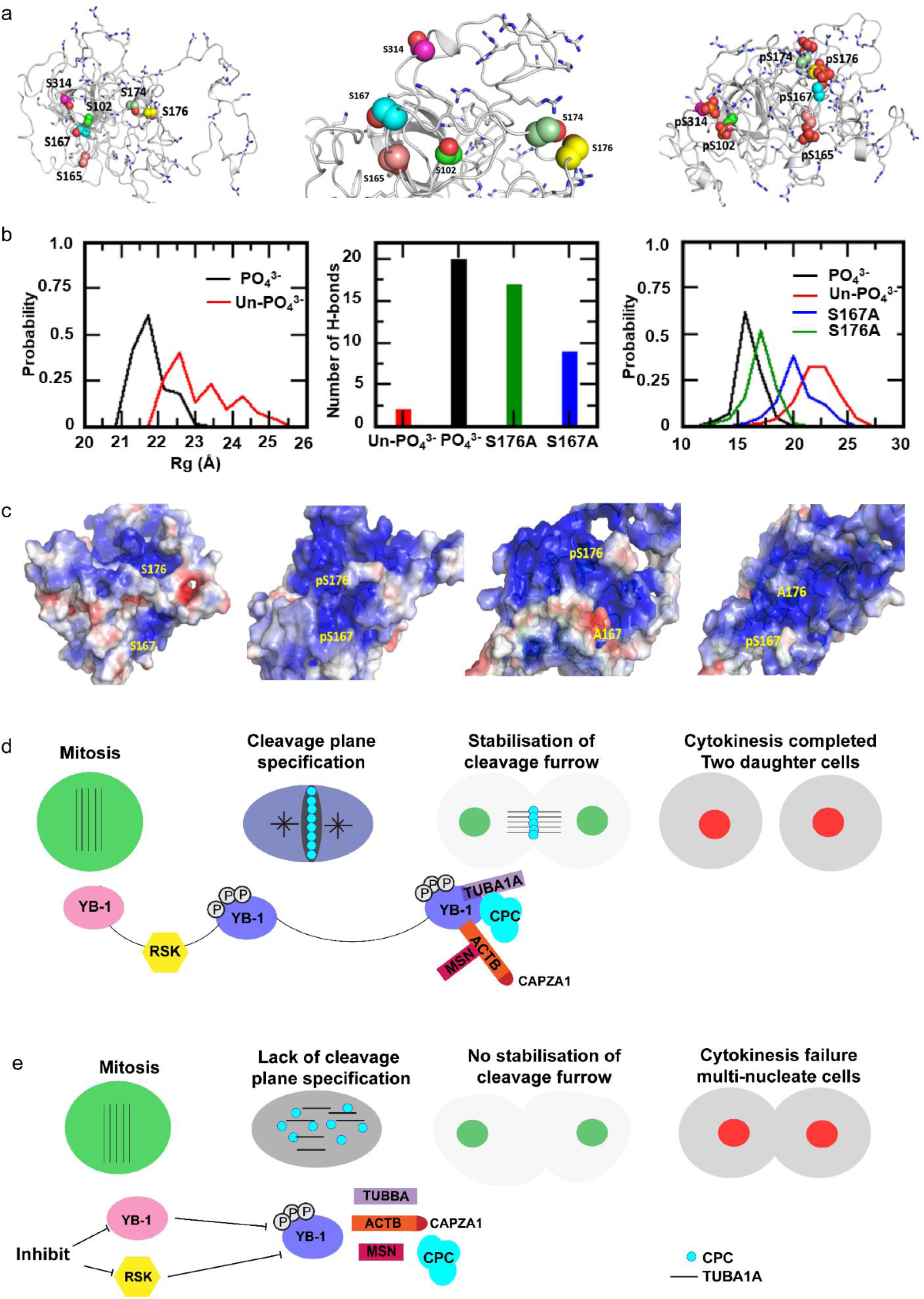
Molecular dynamics simulation models the impact of phosphorylation on localization of the CPC to the cleavage furrow. **a.** YB-1 model generated using ITASSER and MD simulations, left: un-phosphorylated, middle: zoomed view of phosphorylated serines and right: phosphorylated states (pYB-1). Spheres represent serines and phosphoserines, thick grey lines represent positively charged Arg and Lys residues. The ordered Cold Shock Domain is shown at the center flanked by the disordered long termini. **b.** Left: Distribution of the radius of gyration (Rg), a measure of protein compaction. Middle: number of hydrogen bonds involving the serines/phosphorylated serines. Right: distribution of the distances between the backbone atoms of residues 167 and 176 sampled during the MD simulations of YB-1 and phosphorylated YB-1. **c.** Electrostatic surfaces projected on to (left to right) YB1 (unphosphorylated), pYB-1 (phosphorylated), and the two mutants S167A and S176A; the blue and red colours correspond to potentials of +3kcal/mol and -3kcal/mol respectively. **d.** YB-1 has a disordered structure with neutral and negatively charged pockets. Upon phosphorylation by RSK, YB-1 protein undergoes a structural change, creating a positively charged pocket. We speculate that this pocket interacts with the negative charged termini of α-tubulin to define the cleavage plane by facilitating the formation of the microtubule structure and localization of the CPC, which in turn leads to the stabilization of the cleavage furrow. YB-1 also interacts with β-actin and actin related proteins (CAPZA1 and MSN), promoting cytokinesis. **e.** When phosphorylation of YB-1 is inhibited or depleted, the cleavage plane defined by the α-tubulin scaffold cannot form resulting in a failure of CPC localization and cytokinesis.

Folding of YB-1 is maintained by hydrogen bonds formed between the phosphorylated residues and other positively charged amino acids, which is reduced in the absence of phosphorylation (Fig 8b-c). Moreover, the number of hydrogen bonds is greatly reduced when S167 cannot be phosphorylated compared to S176, suggesting that S167 has a greater effect on YB-1 folding compared to S176 (Fig 8b-c). In addition to hydrogen bonds, simulation analyses suggest that phosphorylated YB-1 has a compact structure (lower radius of gyration, Rg) compared to un-phosphorylated YB-1 (Fig 8b-c) and again phosphorylation at S167 has a greater impact on YB-1 folding than S176 (Fig 8b-c). These modelling predications are consistent with our functional data showing that the S167A mutant had a greater negative impact on cytokinesis than the S176A mutant. Cumulatively, the data indicate that RSK phosphorylates YB-1 on multiple residues, thereby altering YB-1 conformation to facilitate CPC recruitment to the cleavage furrow to promote cytokinesis.

## DISCUSSION

YB-1 has long been known to promote cell proliferation [reviewed in ^1^]. Several reports have suggested that YB-1 regulates the G_1_ to S phase transition whilst others have suggested that YB-1 plays a role at G2/M and directly in mitosis ^4,17,22^. However, the precise mechanism is unknown. Using live cell imaging and single cell fate tracking we show that YB-1 is critical for completion of cytokinesis. Cells lacking YB-1 traverse the cell cycle without any defect until cytokinesis, where they fail to undergo cytokinesis, resulting in multinucleation. We further demonstrate that over-expression of YB-1 rescues all these defects. Confocal imaging confirmed that the CPC proteins AURKB, CAPZA1 and MSN failed to localize to the cleavage plane after YB-1 depletion. In addition, using synchronized cells and closely spaced time points, imaging data suggest that YB-1 binds α-tubulin, creating two poles with α-tubulin cables in between where the CPC can bind, defining the position of the cleavage plane and providing a docking site for other proteins required to establish the cleavage furrow. We provide further evidence that zebrafish embryos depleted of *ybx1* fail to assemble and organize the microtubule fibers essential for defining the cleavage plane and completion of cytokinesis. Our results thus provide an explanation for defects in cell proliferation observed in *ybx1*-depleted zebrafish embryos ^16^ and why developmental defects in YB-1 knockout mice are not evident until E11.5 - E13.5 ^14,15^, the time at which there is marked increase in body mass resulting from many cell divisions. Our findings also provide a rationale as to why increased YB-1 levels are often observed in advanced cancers; it likely reflects selection pressures during cancer evolution, assisting in overcoming cytokinesis checkpoints that may be activated during rapid cell division.

YB-1 phosphorylation on S102 is descriptively linked to proliferation and RSK is known to phosphorylate this site ^39,42,46,47^. In our study, in addition to S102, we have identified five highly conserved residues in YB-1 phosphorylated by RSK that are required to facilitate cytokinesis. Mutation of any one of these residues prevented rescue of cytokinesis failure following YB-1 depletion. We also found that these mutants failed to progress through the cell cycle, being ‘stalled’, mainly in G_1_. Using molecular dynamics simulation, we predict that simultaneous phosphorylation of these sites promotes structural alterations in YB-1 protein, resulting in an intense positive potential, which enables YB-1 to interact with the negatively charged termini of α-tubulin ^52^ and actin filaments ^53^. These interactions then allow specification of the cleavage plane and timely localization of the CPC, CAPZA1 and MSN at the cleavage furrow (Fig 8e). YB-1 depletion, or inhibiting YB-1 phosphorylation results in failure to establish the cleavage plane (Fig 8f).

We note there was a continuum of phenotypic severity in cells expressing the YB-1 phosphorylation defective mutants. We propose these differences reflect the impact that each residue has on promoting YB-1 conformational changes following phosphorylation. For example, our modelling data indicate that when S176 and S174 are phosphorylated, they are located in close proximity to a region with a high concentration of positive charges. Phosphorylation of S174 is therefore likely able to compensate when phosphorylation is blocked at S176, but S165 and S167 are located in a region with a lower concentration of positive charges, thus it is unlikely that pS165 is able to effectively compensate for lack of phosphorylation at S167. In agreement, the S167A mutation induced more severe phenotypes than the S176A mutation. Thus, we provide evidence that simultaneous post-translational modification of YB-1 is required to promote its cytokinesis functions.

A previous study ^23^ identified cell division errors following massive over-expression of YB-1 from an inducible promoter. In contrast, we show conclusively through depletion and rescue experiments that YB-1 is a positive regulator that promotes cytokinesis, and in the absence of well-regulated YB-1 function, cytokinesis failure ensues. We anticipate that massive over-expression of YB-1 as used in the above paper, de-regulates the homeostasis within the cytokinesis regulatory network, thereby artefactually inducing cytokinesis failure.

In addition to cytokinesis failure, YB-1 depletion caused an increase in G_1_ transit time of the daughter nuclei of wild type p53 cells that was counteracted by p53 depletion, suggesting the arrest is p53 dependent. p53 activation following cytokinesis failure has been shown to result from activation of the PIDDosome complex or the Hippo pathway in response to supernumerary chromosomes ^54,55^. Both pathways lead to p53 stabilization by inhibiting or degrading MDM2, the E3 ligase that promotes p53 degradation. Thus, it is likely that one or both of these pathways are activated when YB-1 function is impaired. However, with the exception of S176A, the other phosphorylation defective YB-1 mutants exhibited a greater frequency of cells stalled in G_1_ compared with YB-1 shRNA alone. This suggests the G_1_ arrest may not be due solely to the presence of supernumerary chromosomes, but may involve direct interaction of YB-1 with either MDM2 or p53. We anticipate correctly phosphorylated YB-1 can over-ride the checkpoint, possibly by binding directly to p53 ^56^, suggesting that when YB-1 phosphorylation is impaired, the checkpoint remains activated.

In summary, we define precisely how YB-1 regulates cell division. We provide unequivocal evidence that defects in YB-1 result in cytokinesis failure and multinucleation. We demonstrate that YB-1 is essential for microtubule organization and facilitates the specification of the cleavage plane in vitro and in vivo, thus establishing its physiological function. We show that RSK phosphorylates multiple sites on YB-1 and highlight the importance of the YB-1/RSK axis during cytokinesis. Finally, using molecular modeling we demonstrate the importance of structural rearrangement regulated by simultaneous phosphorylation for YB-1 function during cytokinesis. Our work conclusively shows phosphorylated YB-1 is critical for assembly of microtubules, specification of the cleavage plane and localization of the CPC during cytokinesis.

## ACKNOWLEDGEMENTS

This work was supported by grants from the Maurice Wilkins Centre for Molecular Biodiscovery, James Cook Fellowship (JCF-UOO1501), Cancer Society New Zealand (11/07), Health Research Council A*STAR (10-02), Health Research Council of New Zealand (15/500) and the Cancer Institute NSW (05/FRL/1-01). VPM is supported by an Australian Post-Graduate Award from the University of Sydney. SBC acknowledges the Ernest & Piroska Major Foundation for financial support. A.J.C. is supported by grants from the Australian NHMRC (1106241), the Cancer Institute NSW (11/FRL/5-02) and philanthropy from Stanford Brown, Inc (Sydney, Australia). We thank A*STAR and NSCC for computing support. Scott Page and the CMRI ATAC facility supported by the Australian Cancer Research Foundation are thanked for microscopy infrastructure. The WIMR Flow Cytometry Centre supported by the Cancer Council NSW and the Australian NHMRC are thanked for cell sorting. The Otago Zebrafish facility supported by Cure Kids, K D Kirkby Trust, Lottery Health Research NZ and University of Otago are thanked for supporting the Zebrafish work.

## AUTHOR CONTRIBUTIONS

A.B, A.W, A.J.C and S.M conceived ideas and designed experiments, S.M, M.A, A.W, T.A.J, C.M, W.M, K.P, L.H, M.G, P.B, R.H, V.P.M, J.Z and T.K.B performed experiments. S.C and T.K supervised mass spectrometry, M.C assisted with cytokinesis, S.K and C.S.V performed atomistic modeling and A.B, A.J.C and S.M wrote the manuscript.

## DECLARATION OF INTERESTS

SK and CSV are founders/scientific consultants of Sinopsee Therapeutics, a biotechnology company developing molecules for therapeutic purposes; the current work does not conflict with the company. All other authors declare no competing financial interests.

## METHODS

### Cell culture

A549 (human lung adenocarcinoma), HT1080 and HT1080-6TG ^27^ (human fibrosarcoma), MDA-MB-231(triple negative breast cancer) and IIICF/c (skin fibroblast cell line from Li Fraumeni patient ^28^) cells were cultured in Dulbecco’s Modified Eagle Medium (DMEM, Life Technologies or Sigma) supplemented with 10% fetal bovine serum (FBS), 1% Non-essential amino acids (NEAA) and 1% GlutaMax in a 37 °C humidified incubator at 5% CO2 unless stated otherwise. All cell lines were validated for authenticity by CellBank Australia (http://www.cellbankaustralia.com/) using STR profiling. All cell lines are regularly tested negative for mycoplasma contamination.

### Generation of FUCCI cell lines

FUCCI was used to identify cell cycle phases during live cell imaging at a single cell resolution ^57^. mVenus-hGeminin(1/110)/pCSII-EF and mCherry-hCdt1(30/120)/pCSII-EF (a kind gift from Atsushi Miyawaki and Hiroyuki Miyoshi) were individually packaged into lentivectors using the 2nd generation packaging system, and the viral supernatants were used simultaneously to co-infect target cells. Three days post-transduction, cell cultures were sorted at the Westmead Institute for Medical Research (WIMR) flow cytometry core (Sydney, Australia) for mVenus fluorescence, allowed to expand for 5 to 7 days, and sorted again for mCherry fluorescence. Proper progression of red/green coloration during cell cycling was confirmed with live cell imaging as described below before use. FUCCI expressing cell lines used include A549/F, HT1080/F, HT1080-6TG/F and IIICF/c/F.

### siRNA transfection

Cells were reverse transfected with stealth modified 25bp duplex siRNA targeted to YB-1 (si-YB-1 5′-GGUCCUCCACGCAAUUACCAGCAAA-3′) ^4^ and a control siRNA (si-Ctrl 5’ - CCACACGAGUCUUACCAAGUUGCUU-3′) with no known human mRNA targets ^58^. Stealth siRNAs were transfected at a final concentration of 5 nM using Lipofectamine RNAiMax (Invitrogen). Both siRNAs and RNAiMax were diluted in medium without serum. After 10 minutes at room temperature, the diluted RNAiMax was added to the siRNAs, and the mixture was incubated for a further 15 minutes. The lipoplexes formed were added to cells. After overnight transfection, the culture medium was replaced with phenol-free media supplemented with 10% FBS. Media was replaced 24h post transfection on the A549/F, HT1080/F, HT1080-6TG/F, IIICF/c/F and MDA-MB-231 following which the cells were monitored over either 46 hours or 60 hours using the live cell time lapse imaging or were harvested at 48 h for western blot or fixed using 4% PFA for immunofluorescence.

### Generation of YB-1:EBFP2 expression constructs

A two-stage process was used to create HA-tagged YB-1:EBFP2 expression constructs. Firstly an oligonucleotide encoding an HA tag flanked by XhoI and HindIII 5’ and SacII/BamHI 3’ was cloned into the XhoI/BglII sites in the MCS of pEBFP2-N1 plasmid (Adgene 54595) in-frame with EBFP2. The coding sequence of YB-1 and upstream Kozac sequence was amplified from an HA tagged YB-1 plasmid (pHA-YB-1) using TaKaRa Hi Fidelity Taq and cloned upstream of the HA tag XhoI/HindIII. The resulting plasmid produced a YB-1 plasmid tagged with HA and a blue fluorescent protein EBFP2 (YB-1) fusion protein under the control of the CMV promoter. Unique restriction sites within the YB-1 sequence was used to replace wild-type sequence with sequence containing a Serine to Alanine mutation at S102, S165, S167, S174, S176 and S314. All plasmids were propagated through *Stbl3 E. coli* to prevent recombination and sequenced prior to experimental use.

### Plasmid Transfection

Cells (1×10^4^) were seeded in 2ml of DMEM supplemented with 10% FBS in a 12 well plate. After overnight incubation, cells were transfected using Lipofectamine 3000 (LF3000, Invitrogen) with 250ng of shRNA targeting the 3’UTR (sh-YB-1) in combination with 250ng of either Vo:EBFP2 (Vo) or YB-1:EBFP2 (YB-1) or the six phospho-defective mutants (S102A:EBFP2 – S102A, S165A:EBFP2 – S165A, S167A:EBFP2 – S167A, S174A:EBFP2 – S174A, S176A:EBFP2 – S176A and S314A:EBFP2 – S314A). LF3000 (2.5 µl) was diluted in 100 µl of OptiMem, mixed and incubated for 5 min. In parallel, the plasmid was added to 100µl of OptiMem (Invitrogen) along with 2.5µl of P3000 and combined with the tube containing LF3000 and incubated for a further 10 minutes. 200µl of the lipoplexes formed were added to the cells. Media was changed 24 hours post transfection following which the cells were monitored over 60 hours using the live cell time lapse imaging or were harvested at 48 h for western blot as described below.

### Kinase inhibitor treatment and cell cycle

A549/F (1×10^4^) cells were seeded in 1 ml of DMEM supplemented with 10% FBS, 1% NEAA and 1% GlutaMax in a 24 well plate. After overnight incubation media was replaced with media containing either 0.01%(v/v) DMSO or 10nM - 1µM Barasertib (AURKBi), 0.1µM - 5µM BI-D1870 (RSKi), 0.1µM - 100µM MK-2206 HCl (AKTi) and 0.5µM - 50µM Silmitasertib (CSK2i). Confluence was used to measure cell growth by imaging 4 fields per well at 2 hour intervals for 48h using the IncuCyte FLR and accompanying software.

A549/F cells were also seeded (1×10^4^) into 2 ml of DMEM supplemented with 10% FBS, 1% NEAA and 1% GlutaMax in a 12 well plate for live cell time lapse imaging and for western blot. Cells were concurrently seeded (5×10^3^) on glass coverslips in 24 well plates for immunofluorescence. After overnight incubation, DMEM was replaced with medium containing either 0.01%(v/v) DMSO or 1µM Barasertib (AURKB – inhibitor), 5µM BI-D1870 (RSK – inhibitor), 10µM MK-2206 HCl (AKT – inhibitor) and 10µM Silmitasertib (CSK2 – inhibitor). Cells were then either tracked for 46 hours using the live cell time lapse imaging or were fixed at 37°C for 12 minutes with 4% phosphate buffered paraformaldehyde (PFA) at time intervals between 18 and 28 hours following treatment for immunofluorescence.

### Cell extracts and Western Blot

A549/F, HT1080/F, HT1080-6TG/F, IIICF/c/F and MDA-MB-231 (1×10^4^) cells were transiently transfected with either 5nM si-Ctrl, 5nM si-YB-1 a combination of si-YB-1 and si-p53, or Vo or sh-YB-1 as described above. In another experiment, A549 cells were treated with either DMSO or inhibitors of RSK and AURKB. 48 hours post transfection or drug treatment, the media was discarded and the wells were washed with PBS and frozen at -80°C. Cells were lysed in RIPA lysis buffer (150 mM sodium chloride, 1.0% NP-40 or Triton X-100, 0.5% sodium deoxycholate, 0.1% SDS (sodium dodecyl sulfate), 50 mM Tris, pH 8.0). Zebrafish embryos were dechorionated and deyolked prior to lysis. Total protein concentration was determined using the Pierce BCA kit (Thermofisher). Proteins were separated by electrophoresis on Bolt™ 4-12% Bis-Tris Plus Gels (Invitrogen). Proteins were transferred onto nitrocellulose membranes using iBlot gel transfer stacks (Invitrogen). Membranes were blocked with Odyssey® Blocking Buffer (PBS) (Licor) for 1h and then incubated for 1h with the primary antibodies (rabbit anti-YB-1 ^59^ at 1:1000, and mouse anti-β-actin at 1:10,000) diluted in Odyssey® Blocking Buffer (PBS)/0.2% Tween 20 and then incubated for 1h with the secondary antibodies (IRDye® 680RD Goat anti-Mouse IgG (Licor) and IRDye® 800CW Goat anti-Rabbit IgG (Licor)) diluted in Odyssey® Blocking Buffer (PBS)/0.2% Tween 20. Membranes were imaged on the Odyssey® CLx Imaging System (Licor) according to the manufacturer’s instructions. Images were quantified using Image Studio (Licor) software.

### Live imaging and processing of time-lapse data

For live cell imaging, cells were grown in 12-well glass bottom plates (MatTek). Time lapse live cell imaging was performed on a ZEISS Cell Observer inverted wide field microscope, with 20x 0.8 NA air objective, at 37°C, 10% CO_2_ and atmospheric oxygen. Images were captured every six minutes for a duration of sixty hours using an Axiocam 506 monochromatic camera (ZEISS) and Zen software (ZEISS). A ZEISS HXP 120C mercury short-arc lamp and compatible filter cubes were used to obtain fluorescent images and differential interference contrast (DIC) microscopy to capture brightfield images. To achieve optimal image resolution without excessive illumination a binning factor was applied prior to imaging and the ambient conditions were maintained to minimize variations in optical resolution and illumination. The acquired videos were analyzed using the Zen Blue software (Zeiss). For all videos, mitotic duration and outcomes were scored by eye and calculated from nuclear envelope breakdown until cytokinesis. FUCCI videos were scored by eye, for G1 (red) and S/G2/M (green). During time lapse imaging, some cells migrated away from the field of view, so the fate of those cells could not be determined and they were removed from further analyses.

Time lapse live cell imaging was also carried out on MDA-MB-231 cells that were transiently transfected with siRNA, seeded into a 2-well glass bottom µ-slide (ibidi) and subsequently cultured in a heated and humidified chamber (ibidi) under cell culture conditions (37°C, 5% CO_2_). Images were acquired every 15 minutes for 48 hours following transfection using a Nikon Ti-E inverted microscope with a 20× 0.75 NA dry objective and equipped with a Nikon DSRi2 CCD camera. The acquired images were analyzed with NIS elements (Nikon) and ImageJ software.

### Cell synchronization

A549 (1×10^4^) cells were seeded on 13 mm diameter coverslips in a 24 well plate. After overnight incubation, cells were treated with 2mM thymidine (Sigma) for 18h, washed twice with PBS and released into medium without thymidine for 7 – 8 hours. The cells were again treated with 2mM thymidine (Sigma) for 15 – 16 hours, washed twice with PBS and incubated in medium without thymidine for 6-7 hours following which cells were fixed in 4% phosphate buffered paraformaldehyde (4% PFA) for 12 minutes and then washed thoroughly with PBS.

### Cell proliferation assays

A549 (2×10^3^) cells were seeded in 80µl of DMEM supplemented with 10% FBS in a 96 well plate. After overnight incubation, 80µl of media containing either 0.01%(v/v) DMSO or 1µM Barasertib (AURKB - inhibitor) or 5µM BI-D1870 (RSK - inhibitor) was added to the cells. In parallel, cells were also transfected with 25ng of either Vo or YB-1:EBFP2 using Lipofectamine 3000 (LF3000, Invitrogen) as described above. Each treatment had six biological replicates. 20µl of the lipoplexes were added to the cells. At the indicated time points the medium was removed and the plates were frozen at - 80°C. After all the time points had been collected the plates were thawed and the DNA content measured using a SYBR Green I-based fluorimetric assay as described previously ^58^. Briefly, 100 µl of lysis buffer (10mM Tris-HCl pH 8.0, 2.5mM EDTA and 1%(vol/vol) Triton X-100) containing 1:4000 SYBR Green I (Invitrogen) was added to the wells and the plates were incubated overnight at 4°C. The plates were mixed and the fluorescence signal for each well was measured for 1 second at an excitation of 485nm and emission of 535nm using a BioTek microplate reader (BioTek). Growth curves were plotted as the fluorescence values at each time point.

### Immunofluorescence

Fixed cells were thoroughly washed in PBS, quenched with 0.1M glycine in PBS, permeabilized with 0.2% Triton X-100 (SigmaAldrich) and following further washing with PBS, were blocked with 1% BSA in PBS with 0.2% cold fish skin gelatin (SigmaAldrich) prior to incubation with primary antibodies in blocking solution. Following primary incubation, cells were again washed thoroughly to remove any unbound primary antibody and then incubated with a fluorophore-labelled secondary antibody. Sequential immunolabelling was carried out in some instances, whereby cells were thoroughly washed with 0.1%TWEEN-20 (SigmaAldrich) in PBS and re-blocked prior to incubation with another primary. Filamentous (F)-actin was labelled with directly conjugated Alexa Fluor 568 Phalloidin (ThermoFisher Scientific). The DNA-binding dye Hoescht 33258 (Molecular Probes) was used to visualise cell nuclei. Cells were thoroughly washed in PBS prior to mounting in Fluoromount-G (SouthernBiotech) on glass slides.

### Antibodies

The following primary antibodies were used: rabbit anti-YB-1 polyclonal to N-terminal (generated in house ^59^ 1:1000), sheep anti-YB-1 polyclonal to N-terminal (generated in house), mouse anti-AIM-1 (611082, BD transduction laboratories, 1:200); mouse anti-α-Tubulin (DM1A, 3873, Cell Signaling Technology, 1:2000) mouse anti-CAPZA1 (66066-1, proteintech 1:200); mouse anti-Moesin (MSN/491, Novus biologicals, 1:200). Secondaries used were goat anti-rabbit Alexa Fluor 488 (Molecular Probes, 1:1000), F(ab’)2 goat anti-mouse Alexa Fluor 350 (Molecular Probes 1:1000) and F(ab’)2 goat anti-mouse Alexa Fluor 633 (Molecular Probes, 1:1000).

### Zebrafish lines

Wildtype WIK zebrafish (**ZFIN ID:** ZDB-GENO-010531-2) were maintained as described previously ^60^. Stages were determined by using both time since fertilisation (hpf) and morphological features ^61^

### Microinjection of zebrafish embryos

Antisense morpholino oligonucleotides were obtained from GeneTools LLC. For microinjection, 1 nl of morpholino solution (4µg/µl) diluted in Danieau’s buffer was injected into the yolk of embryos at the 1 to 2-cell stage. Morpholino sequences were MO_*ybx1* (morphilino targeting ybx1) 5′-GTTGTGTCTCGGCCTCGCTGCTCAT-3′ and MO_Ctrl (control morpholino), 3′-TAATTTACTTACCCTCAAGTTGCTG-5’.

### Whole-mount immunofluorescence of zebrafish embryos

Whole-mount immunofluorescence was carried out according to a standard protocol, as previously described ^62^, In brief, embryos were fixed in 4% PFA in PBS at 4°C overnight, dechorionated and stored in methanol at -20°C. Following sequential rehydration into PBS with 0.1% Tween (PBSTw), embryos were immersed overnight in 30% sucrose at 4°C before incubation at 70°C in 150 mM Tris-HCl (pH 9.0). Following thorough washing with PBSTw, they were then blocked with 10% sheep serum, 0.8% Triton, 1% BSA in PBSTw for 3 hours at 4°C before incubation with primary antibodies in 1% sheep serum, 0.8% Triton, 1% BSA in PBSTw for 3 days at 4°C on a rotating mixer. Residual primary antibody was then removed by successive 1hr washes in PBS with 10% sheep serum and 1% Triton before incubation in secondary antibodies and Hoechst dye in the dark for 2.5 days. Following removal of residual secondary antibodies by successive 1hr washes in PBS with 10% sheep serum and 1% Triton, embryos were mounted in Fluoromount-G (SouthernBiotech) and imaged on a Nikon A1 inverted confocal microscope. Antibodies used were rabbit anti-YB-1 polyclonal to N-terminal (generated in house ^59^ 1:1000), mouse anti-α-Tubulin (DM1A, 3873, Cell Signaling Technology, 1:500). Secondary antibodies used were F(ab’)_2_ goat anti-rabbit Alexa Fluor 568 (Molecular Probes, 1:1000), goat anti-mouse Alexa Fluor 488 (Molecular Probes 1:1000).

### Subcellular fractionation of cultured cells

Cells were enriched into subcellular fractions as described previously ^59^. Briefly, 1 × 10^4^ cells per µl were swelled in a hypotonic buffer (10 mM HEPES, pH 7.9, 1.5 mM MgCl_2_, 10 mM KCl, 0.5 mM, DTT, 1 × Complete EDTA-free), and incubated for 5 min at 4°C before being transferred to a Dounce homogeniser (Kontes, 885300-0002, 2 ml, tight pestle; 0.013 - 0.064 mm clearance). The Dounce was applied until 95% of the cell membranes were ruptured and lysate was spun at 220 g for 5 min at 4°C. The supernatant was collected and stored as cytoplasmic extract. The pelleted nuclei were re-suspended in 5 ml of chilled low sucrose solution (0.25 M Sucrose,10 mM MgCl_2_) which was gently layered over 5 ml of chilled high sucrose solution (0.88 M Sucrose,10 mM MgCl_2_). The nuclei were spun at 2800 g for 10 min at 4°C to remove contaminating cytoplasmic proteins.

### Immunopurification of YB-1

YB-1 was immunopurified as described previously by Cohen et al ^59^. Lysates were prepared at the equivalent of 1 × 10^4^ cells per µl for whole cells or cytoplasmic lysates and 5 × 10^4^ cells per µl for nuclear lysates in a low detergent RIPA buffer (50 mM Tris-HCL, 150 mM NaCl, 0.5% NP40, 0.1% deoxycholate). Complete lysis of nuclei was ensured by passing the nuclei pellets and lysis buffer through a 22 gauge needle 10 times followed by incubation for 20 min at 4°C. Debris was cleared from lysates by centrifugation at 13000 g for 10 min. All lysates included protease and phosphatase inhibitors at the recommended concentration (Complete-EDTA-free and PhosSTOP [Roche]).

Protein G dynabeads were prepared for use following the manufacturer’s instructions (Life Technologies). Protein G dynabeads were immobilized on a magnet and washed three times with 300 µl of low-detergent RIPA buffer. 100 µg of anti-YB-1 was added to the lysates and incubated with gentle agitation for 1 hr at 4°C. Twenty-five µl of protein G beads were then added for every 20 µg of antibody and incubated for another 30 min at 4°C. The bead:antibody:YB-1 complexes were then immobilized by a magnet, the lysate removed, and the interaction of YB-1 and the anti-NYB1 was disrupted by the addition of the peptide to which NYB1 was raised at a concentration of 100-fold compared to the antibody. The immunopurified proteins were then stored at -80°C to await further analysis.

### Mass spectrometry

For detection of phosphorylated YB-1 peptides, phospho-enriched peptide fractions were prepared from whole cells. Peptides were generated from 500 µg of protein from whole cell lysates by tryptic protein digestion using the filter-aided sample preparation method ^63^. Phosphorylated peptides were enriched from 450 µg of digested protein peptides using a TiO2/ZiO2 TopTip following the manufacturers protocol (TT2TIZR, Glygen). YB-1 enriched phosphorylated peptides were re-solubilised in 2% (v/v) acetonitrile, 0.2% (v/v) formic acid in ultrapure H_2_O and separated using an LC-coupled TripleTOF 5600 + (AB Sciex). Peptides were separated on emitter-tip columns (75 µm ID fused silica tubing packed with C-18 material on a length of 12 cm). The gradient for liquid chromatography was modified depending on the complexity of the fraction or band. Generally, the gradient developed from 1% [v/v] acetonitrile, 0.2% (v/v) formic acid to 80% (v/v) acetonitrile, 0.2% (v/v) formic acid in ultrapure H_2_O at a flow rate of 400 nl/min.

Full MS (MS1) in a mass range between m/z 400 - 2000 was performed in the Orbitrap mass analyzer with a resolution of 60,000 at m/z 400 and an AGC target of 2e5. The strongest 5 signals, with a charge state ≥ [M + 2H]2+, were selected for collision induced dissociation -MS/MS (MS2) in the LTQ ion trap at a normalized collision energy of 35% using an AGC target of 1e5 and two microscans. Dynamic exclusion was enabled with 2 repeat counts during 30 second (sec) and an exclusion period of 180 secs. The exclusion mass width was set to 0.01.

Phosphorylated YB-1 peptides using targeted mass were detected using multi-reaction monitoring mode where the precursor m/z of phosphorylated YB-1 peptides that were detected in immunopurified YB-1 samples were subjected to MS2 scans throughout the gradient. Levels of YB-1 in the enriched phosphorylated samples were also estimated by using a similar gradient and sequential window acquisition of all theoretical fragment ion spectra mass spectrometry (SWATH) method on a TripleTOF 5600 + mass spectrometer. The data from targeted LC-MS/MS and SWATH were analysed using Mascot (http://www.matrixscience.com) and Skyline (v4.2.0). Quantification of phosphorylation at different sites on YB-1 was performed using the peak areas from fragment ions unique to individual phosphorylation sites.

### Modeling the structure of YB-1 protein

The automated I-TASSER pipeline ^49^ which generates atomic models of proteins based on multi-threading alignment and iterative template fragment assembly simulations was used. This yielded several models of which we chose the model that incorporated the coordinates of the CSD of YB-1 whose 3-dimensional structure has been resolved by NMR ^50^ and was one of the templates used by I-TASSER.

### Molecular dynamics simulations

The 3-dimensional model of full length YB-1 was subject to molecular dynamics (MD) simulations in its phosphorylated form (at Serines 102, 165, 167, 174, 176 and 314, referred to as pYB-1), unphosphorylated form and of mutant forms, S167A and S176A (where the two mutant forms had the other 5 serines phosphorylated) using the *pemed.CUDA* module of the program Amber14 ^64^. The all atom version of the Amber 14SB force field (ff14SB) ^65^ was used for the protein. Force field parameters for phosphorylated serines were taken as described elsewhere ^66^. For the phosphate groups, an overall charge of -2e was used. The *Xleap* module of Amber14 was used to prepare the system for the MD simulations. All the simulation systems were neutralized with appropriate numbers of counterions. Each neutralized system was solvated in an octahedral box with TIP3P ^67^ water molecules, leaving at least 10 Å between the solute atoms and the borders of the box. All MD simulations were carried out at 300K. During the simulations, the long-range electrostatic interactions were treated with the particle mesh Ewald ^68^ method using a real space cutoff distance of 9 Å. The Settle ^69^ algorithm was used to constrain bond vibrations involving hydrogen atoms, which allowed a time step of 2 fs during the simulations. Solvent molecules and counterions were initially relaxed using energy minimization with restraints on the protein. This was followed by unrestrained energy minimization to remove any steric clashes. Subsequently the system was gradually heated from 0 to 300 K using MD simulations with positional restraints (force constant: 50 kcal mol^-1^ Å^-2^) on the protein over a period of 0.25 ns allowing water molecules and ions to move freely. During an additional 0.25 ns, the positional restraints were gradually reduced followed by a 2 ns unrestrained MD simulation to equilibrate all the atoms. For each system, three independent MD simulations (assigning different initial velocities) were carried out for 250 ns with conformations saved every 10 ps. Simulation trajectories were visualized using VMD ^70^ and figures were generated using Pymol ^71^.

