## Supplementary Fig 1, Supplementary Fig 2, Supplementary Fig 3, Supplementary Fig 4 for "Critical role for cold shock protein YB-1 in cytokinesis"

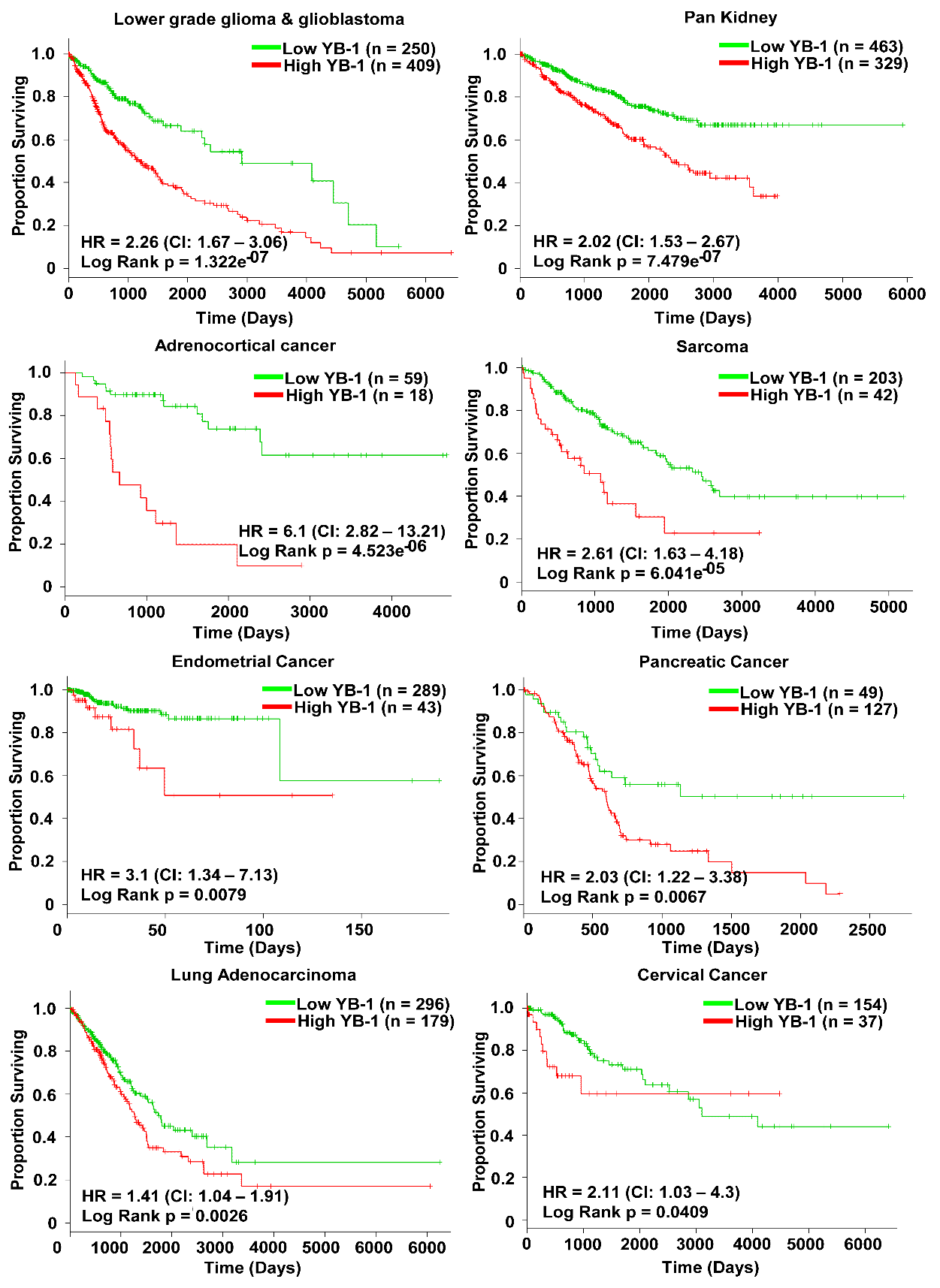

**Supplementary Figure 1: Significant prognostic value of *YBX1* mRNA in cancer.** Kaplan-Meier plot of high risk versus low risk tumors stratified by *YBX1* mRNA expression from TCGA. Patients with high risk were identified using the algorithm by SurvExpress which performs a log-rank test along all values of the ordered prognostic index and then chooses the split point where the p value is minimum. Patients in the high risk group had high expression of *YBX1* mRNA (red) in their tumors compared to those the patients in low risk group (green). All analyses were carried out using the SurvExpress web resource.

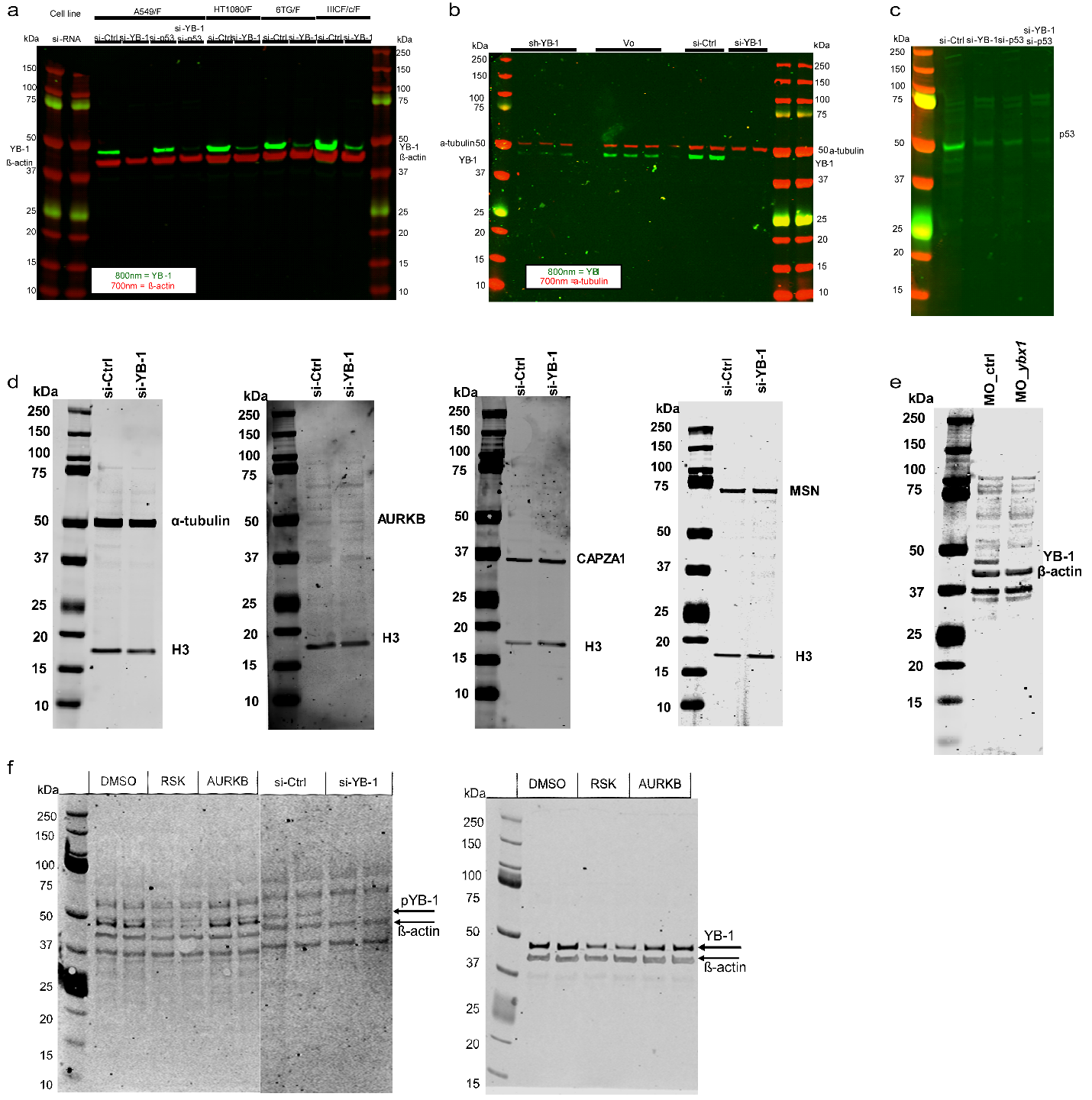

**Supplementary Figure 2: Western Blots. a.** Western blot of YB-1 protein after treating cells for 48h with either si-YB-1 (5nM) or si-Ctrl (5nM) in A549/F, HT1080/F, HT1080-6TG/F and IIICF/c/F cells. **b.** Western blot of YB-1 protein after treating cells with 48h of either si-YB-1 (5nM) or si-Ctrl (5nM) in MDA-MB-231 cells or treating A549/F cells with either an sh-YB-1 targeting the 3’UTR of YB-1 (1µg) or Vo (1µg). **c.**Western Blot of p53 knockdown in A549/F cells after treating cells with either si-Ctrl, si-YB-1 or si-p53 for 48 hours. **d.** Western blot showing the levels of α-tubulin, AUKRB, CAPZA1, MSN in A549/F cells treated for 48h with either si-YB-1 (5nM) or si-Ctrl (5nM). **e.** Western blot of *ybx1* protein after treating zebrafish embryos with either MO_Ctrl or MO_*ybx1*. **f.** **Left:** Western blot of phosphorylated YB-1 at S102 in A549 cells treated with either DMSO, RSK inhibitor, AURKB inhibitor, si-Ctrl or si-YB-1. **Right:** Western blot of YB-1 protein in cells treated with either DMSO, BI-D1870-5µM (RSK) or Barasertib-1µM (AURKB).

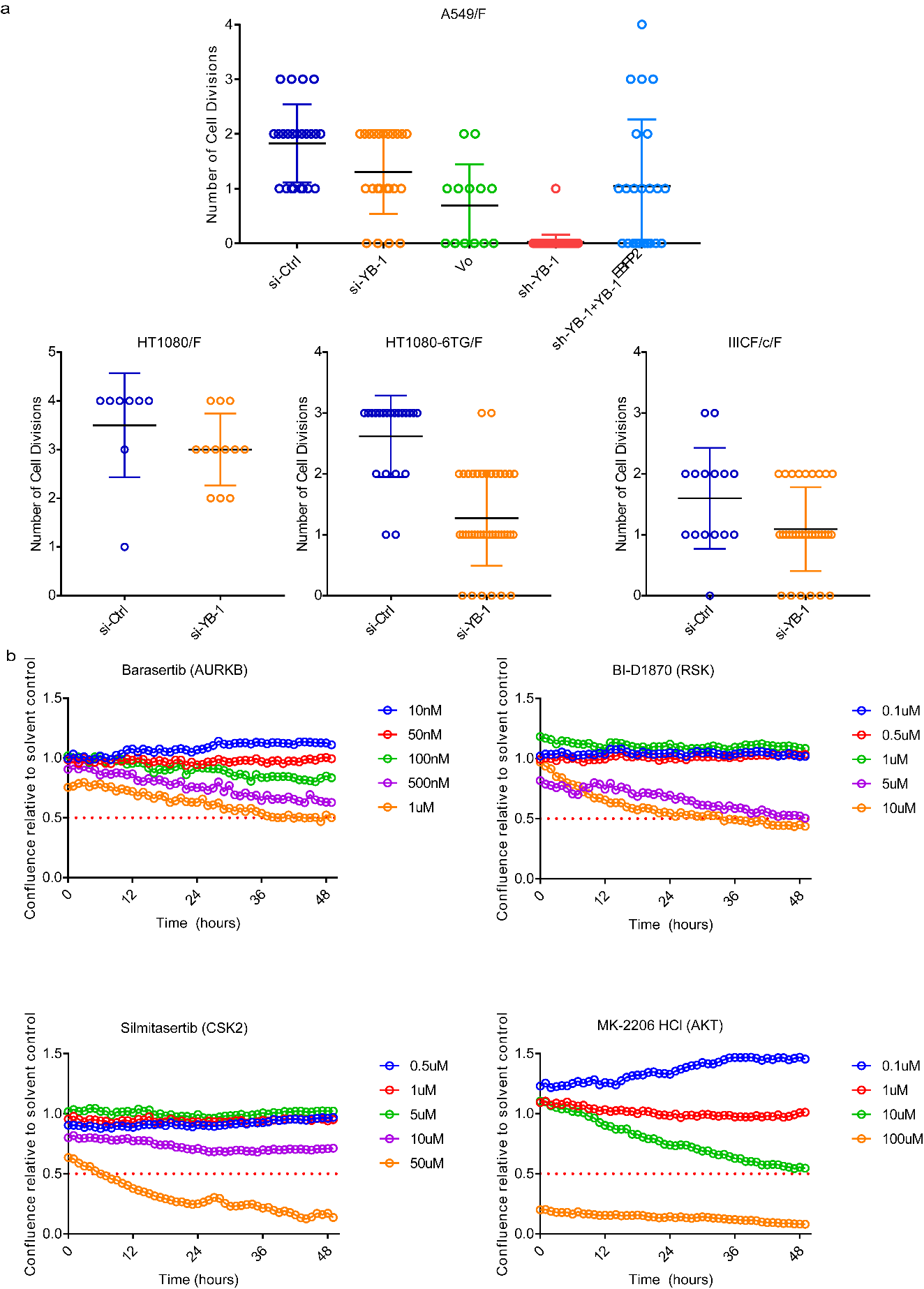

**Supplementary Figure 3: a. YB-1 depletion decreases number of cell divisions.** Number of cell divisions of A549/F cells in cultures treated with si-Ctrl, si-YB-1, Vo, sh-YB-1 and sh-YB-1+YB-1^BFP2^ post transfection (top panel).Number of cell divisions of HT1080/F, HT1080-6TG/F and IIICF/c/F cells treated with either si-Ctrl or si-YB-1 post transfection (bottom panel). The line in the middle of each box represents the median, the top and bottom outlines of the box represent the first and third quartiles. Significance was determined using Mann-Whitney U test, * p < 0.05, *** p < 0.001**. b.** **Dose dependent decrease in growth of A549/F cells treated with kinase inhibitors.** Growth response of A549/F cells treated with: 10nM - 1µM Barasertib (AURKB - inhibitor), 0.1µM - 10µM BI-D1870 (RSK - inhibitor), 0.1µM - 100µM MK-2206 HCl (AKT - inhibitor) and 0.5µM - 50µM Silmitasertib (CK2 - inhibitor). Confluence was used to measure cell growth by imaging 4 fields per well at 2 hour intervals for 48h using the IncuCyte FLR and accompanying software.

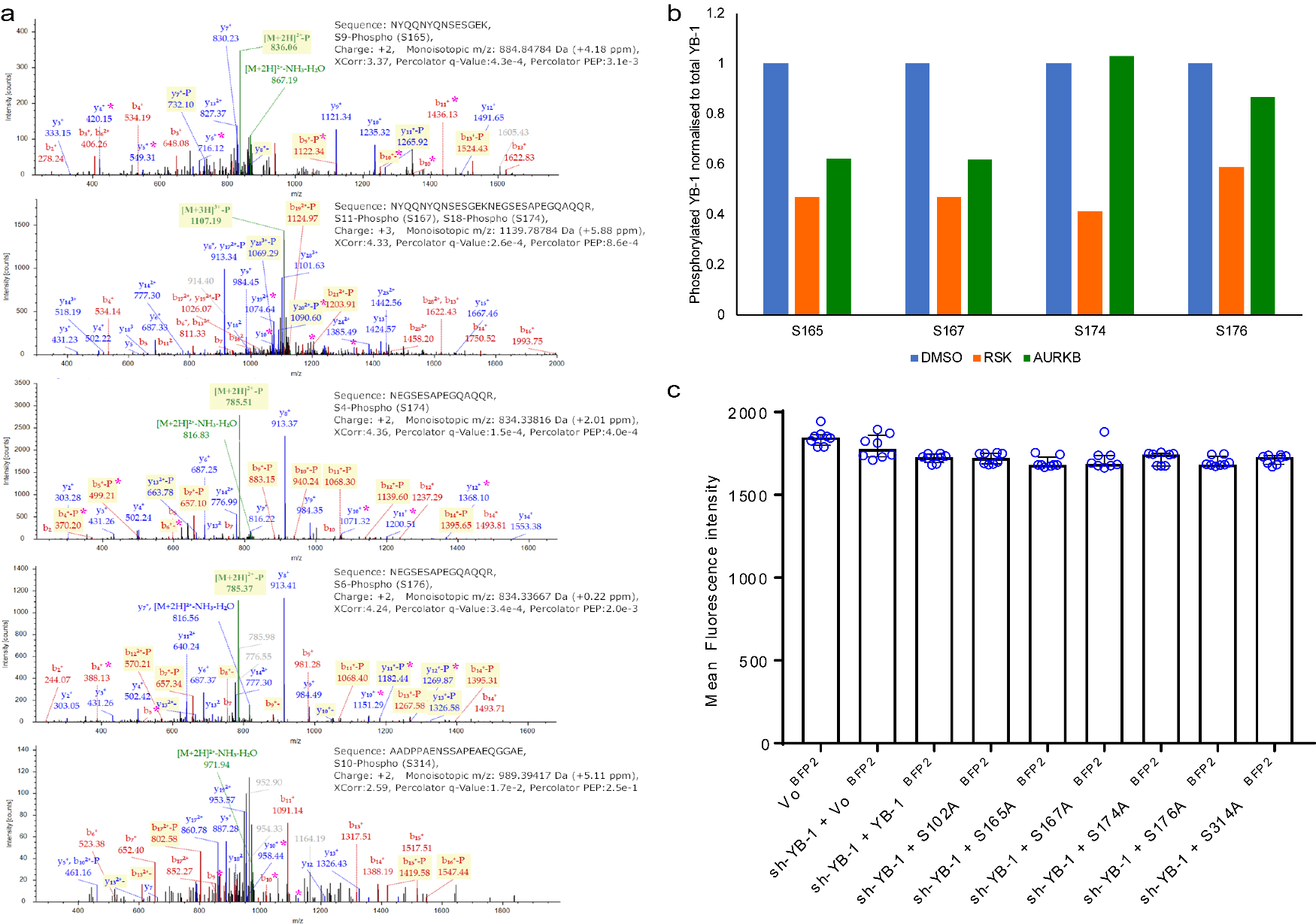

**Supplementary Figure 4: a. Representative MS/MS spectra showing phosphorylation of YB-1 at residues S165, S174, S176 and S314.** The annotated peaks were identified as b-ions (red) and y-ions (blue) by Sequest. Phosphorylation was identified as the addition of HPO_3_ (+ 79.96633 Da) or the neutral loss of H_3_PO_4_ (- 97.976896; annotated by “-P” and a yellow background). Ions produced by neutral losses from the precursor ion are shown in green. Peaks that clearly indicate the site of the phosphorylation are annotated with a “*”. **b. RSK inhibition prevents phosphorylation of YB-1 at multiple serines**. Mass spectrometry analyses of phosphorylated YB-1 at S165, S167, S174 and S176 in A549 cells treated with either DMSO, Barasertib-1µM (AURKB) or BI-D1870-5µM (RSK). **c. EBFP2 constructs expressed similar levels of protein.** The graph shows the average of the arithmetic mean intensity of the individual EBFP2 constructs determined by time lapse live cell imaging. The bars and error bars represent the mean ± s.d. across eight imaging fields.
